# Model-based evaluation of critical nursery habitats for juvenile blue crabs through ontogeny: abundance and survival in seagrass, salt marsh, and unstructured bottom

**DOI:** 10.1101/2023.07.20.549877

**Authors:** A. Challen Hyman, Grace S. Chiu, Michael S. Seebo, Alison Smith, Gabrielle G. Saluta, Kathleen E. Knick, Romuald N. Lipcius

**Affiliations:** Virginia Institute of Marine Science, William & Mary, Gloucester Point, VA 23062, USA; Australian National University; Virginia Commonwealth University; University of Washington; University of Waterloo

**Keywords:** *Callinectes sapidus*, larval supply, ecosystem based fishery management.

## Abstract

Nursery habitats confer higher food availability and reduced predation risk, thereby contributing disproportionately more individuals per unit area to the population compared with other habitats. Nursery status is inferred through evaluation of four metrics: density, growth, survival, and linkage between juveniles and adults. Moreover, organisms commonly use multiple habitats as nurseries throughout ontogeny to satisfy shifting resource requirements. To this end, we conducted manipulative and mensurative field experiments to evaluate two nursery metrics, abundance and survival, for juvenile blue crabs across multiple size classes and habitats, including structurally complex habitats — seagrass meadows and salt marshes — and unstructured habitat (sand flats) in the York River, Chesapeake Bay. We also considered effects of site-specific spatial orientation within the York River, seasonality, physicochemical variables, and postlarval influx. Our results showed that abundance was higher in both seagrass meadows and salt marshes relative to unstructured sand, and positively associated with turbidity and post-larval abundance. Notably, seagrass habitats harbored the highest abundances of small (*≤* 15 mm carapace width) juveniles, whereas salt marsh edge harbored the highest abundance of medium (16–30 mm carapace width) and large (31– 60 mm carapace width) juveniles. Moreover, survival was positively associated with juvenile size and structurally complex habitats relative to unvegetated controls. Seasonally, survival peaked in April, reached a seasonal minimum in August, and increased throughout fall. Finally, habitat-specific survival was dependent on spatial position: survival was elevated at upriver salt marsh and unstructured sand habitats compared to downriver counterparts. In nursery habitats of marine species, evaluation of survival, growth, abundance and ontogenetic habitat shifts has typically focused on relatively broad size ranges through the juvenile phase. Yet, ontogenetic shifts in habitat use may occur within much narrower size ranges, which has not been well studied and which is critical to the conservation and restoration of nursery habitats. We found that habitat-specific utilization rates differed by juvenile size class over a surprisingly narrow range of size, and were related to (1) the structural and biological characteristics of the nominal nursery habitats, (2) spatial gradients of environmental variables within the tributary, and (3) the likely trade-offs between growth and survival through ontogeny. Taken together, abundance and survival results indicate that seagrass meadows are key nurseries primarily for early-stage juveniles, whereas salt marshes are an intermediate nursery habitat for larger individuals to maximize growth-to-mortality ratios. Our results underscore the need to consider both habitats as critical nurseries for juvenile blue crabs throughout ontogeny.

## 1. Introduction

Estuaries are highly productive ecosystems that sustain populations of many commercially exploited marine and estuarine species. Productivity within estuaries is driven by shallow-water habitats such as seagrass beds, salt marshes, and mangrove forests. These habitats serve as valuable nurseries for commercially exploited fish and invertebrate species (Beck et al., 2001; Minello et al., 2003; Heck Jr, Hays and Orth, 2003; Litvin et al., 2018; Lefcheck et al., 2019). Nursery habitats are characterized by abundant food and refugia, and thus provide conditions favorable for juvenile growth and survival. Abundant and diverse nursery habitats elevate secondary production (Hyman et al., 2022), while loss of nursery habitat may cause population declines even when fishing pressure is low (Deriso, Maunder and Pearson, 2008; Vasconcelos et al., 2014). Identifying and understanding nursery functions and contributions for ecologically and economically important species has therefore been a major focus of marine research (Beck et al., 2001; Heck Jr, Hays and Orth, 2003; Minello et al., 2003; Nagelkerken et al., 2015; Litvin et al., 2018; NMFS, 2010; Peters et al., 2018).

The value of a particular habitat to juveniles is dependent on size- or life-stage due to changing growth and survival requirements (Werner and Gilliam, 1984; Dahlgren and Eggleston, 2000; Lipcius et al., 2005; Fodrie, Levin and Lucas, 2009). Juveniles possess life-history strategies that maximize energy gains (i.e. growth rates) and minimize predation risk (Werner and Gilliam, 1984; Dahlgren and Eggleston, 2000). In many cases, however, there are trade-offs; habitats that offer higher potential growth rates can possess greater risks of predation (Lipcius et al., 2005; Seitz, Lipcius and Seebo, 2005). Juvenile habitat utilization therefore shifts with transitions between different life stages because of changing resource needs as well as altered predation risk. Initially, the earliest juvenile life stages are the smallest and most vulnerable to predation, and typically prioritize refuge for survival over food availability for growth (Johnston and Lipcius, 2012). As juveniles grow, their probability of survival increases as their size exceeds the mouth gapes of many smaller predators (Nakamura et al., 2012). However, to continue growth and development juveniles must have access to ample food resources. Consequently, juveniles initially prioritize structure over food availability, but will increasingly seek out habitats with higher food supplies as they continue to develop.

Literature on nursery habitats has increasingly emphasized the need to consider all critical habitats utilized by a species through ontogeny (Nagelkerken et al., 2015). Prioritizing only a subset of nursery habitats throughout ontogeny may miss bottlenecks at one or more life stages. For example, if initial settlement habitats are conserved, but intermediate habitats used by larger juveniles are allowed to deteriorate, secondary production of juveniles will decrease. Explicitly focusing on vital rates for only a single size class or multiple size classes in aggregate are insufficient for complete nursery inference (Sheaves, Baker and Johnston, 2006; Sheaves et al., 2015); multiple size classes must be evaluated concomitantly across candidate habitats to identify the full scope of nursery habitats required to maintain healthy populations (Nagelkerken et al., 2015; Hyman et al., 2023).

The blue crab *Callinectes sapidus* is a commercially exploited, estuarine dependent species which exhibits ontogenetic changes in habitat utilization (Lipcius et al., 2007). Similar to many estuarine-dependent species with complex, bi-phasic life histories, blue crabs experience multiple larval, post-larval, and juvenile phases with varying requirements prior to reaching maturity. Following advection to the continental shelf for growth and development of larval stages, postlarvae (herein megalopae) re-enter estuaries and settle into nursery habitats (Epifanio, 2007, 2019). The current paradigm of blue crab life history maintains that megalopae primarily seek out structurally complex habitats due to their refuge quality (Epifanio, 1995; van Montfrans, Ryer and Orth, 2003; Epifanio, 2007, 2019). Elevated abundance and survival of juvenile blue crabs in seagrass beds relative to unstructured habitats led multiple investigators to conclude that seagrass beds were the preferred initial nursery habitat (Orth and van Montfrans, 1987; Lipcius et al., 2005; Ralph et al., 2013). In regions where seagrass beds were locally sparse or absent from the estuarine habitat mosaic, blue crabs opportunistically exploited salt marshes, macroalgae, and other alternative nurseries (Wilson, Able and Heck Jr, 1990a,b; Thomas, Zimmerman and Minello, 1990;

Posey et al., 2005; Johnson and Eggleston, 2010; Rudershausen, Merrell and Buckel, 2021). Although juvenile survival is higher in structured habitats, juvenile growth rates appear higher in turbid marsh-fringed habitats distant from SAV beds (Seitz et al., 2003; Seitz, Lipcius and Seebo, 2005; Lipcius et al., 2005). Based upon mechanisms governing ontogenetic habitat shifts (Werner and Gilliam, 1984), previous studies proposed juvenile blue crabs remain in structurally complex habitats such as seagrass beds until reaching an initial size refuge from predation at 20 – 30 mm carapace width (CW), upon which they disperse to unstructured habitats that minimize mortality-to-growth ratios (Lipcius et al., 2007; Johnston and Lipcius, 2012).

Recent evidence considering multiple size classes of small juveniles indicates that certain structurally complex habitats may be used differently by varying juvenile blue crab size classes before dispersing to unstructured habitat. Specifically, abundances of small (*≤*15 mm CW) juveniles are consistently highest in seagrass beds (e.g. Hyman et al., 2023) partially due to the relatively high structural complexity associated with seagrass shoots (Lipcius et al., 2007). Seagrass structure enhanced survival of small juveniles in controlled mesocosms (Orth and van Montfrans, 2002) and in field experiments (Pile et al., 1996; Hovel and Lipcius, 2001, 2002; Bromilow and Lipcius, 2017). However, this benefit may diminish at larger size classes. Lower juvenile predation rates in mesocosms suggest that juvenile blue crabs achieve a size refuge from smaller predators at sizes as small as 12 mm CW (Orth and van Montfrans, 2002). Meanwhile, abundances of larger (i.e. *>*15 mm CW) size classes are higher in salt marshes (Ralph, 2014; Rudershausen, Merrell and Buckel, 2021; Hyman et al., 2022, 2023). Although less structurally complex than seagrass beds, salt marsh shoot density impedes foraging success of subadult and adult blue crab predators on conspecific juveniles in predation experiments (Johnston and Caretti, 2017; Miller et al., 2023). Tidal marsh creeks – particularly in turbid conditions – also support high densities of thin shelled bivalves preferentially consumed by larger juveniles (Seitz et al., 2003; Seitz, Lipcius and Seebo, 2005). Collectively with habitat- and size-specific patterns in juvenile abundance, these observations suggest that salt marshes represent an intermediate nursery habitat following initial dispersal from seagrass beds but before utilization of unstructured habitat (Ralph, 2014; Hyman et al., 2023), potentially due to changes in mortality-to-growth ratios.

This recent evidence of differential early-life habitat utilization among size classes indicates that the current paradigm of blue crab early life history requires revision. Although previous mesocosm experiments suggested juvenile blue crab survival is higher in salt marshes relative to unstructured habitats (Johnston and Caretti, 2017; Miller et al., 2023), there is little field evidence to validate this hypothesis. For example, juvenile survival was equivalent in salt marshes and unstructured habitat in a fragmented salt marsh system in the Gulf of Mexico (Shakeri et al., 2020), and no comparable field studies exist to evaluate juvenile survival among multiple structured habitats in mid-Atlantic estuaries. As habitat-specific secondary production is a function of both abundance and survival, these vital rates should be assessed jointly to understand the relative importance of initial nursery habitats and intermediate habitats within the estuarine seascape for a given life stage (Beck et al., 2001; Nagelkerken et al., 2015).

In this study, we conducted mensurative (abundance, postlarval influx) and manipulative (survival) field experiments to investigate juvenile blue crab abundance and survival across multiple nursery habitats and juvenile size classes. Our objectives were to 1) evaluate the extent to which habitat- and size-specific vital rates were consistent with the existing paradigm on juvenile blue crab life history and 2) determine the contribution of different structurally complex habitats to juvenile blue crab secondary production for different juvenile stages. Specifically, we constructed Bayesian hierarchical models to evaluate the effects of two structurally complex habitats – seagrass beds (herein, seagrass) and salt marsh edge (herein, SME) – as well as unstructured sand habitat (as a control; herein, sand) across the seascape of the York River, a tributary of Chesapeake Bay. We focused on three size classes of juveniles: small (*≤*15 mm CW), medium (16–30 mm CW) and large (31–60 mm CW). In addition, we assessed the influence of spatially-varying turbidity and megalopal supply through complementary field sampling.

## 2. Methods

### 2.1. Study Area

Blue crab abundance sampling and survival experiments were conducted in the York River, a tributary in the lower portion of western Chesapeake Bay. The tributary contains a wide range of habitat configurations and gradients of environmental variables such as turbidity (Hyman et al., 2023). In addition, the York River harbors high abundance of juvenile blue crabs spanning multiple size classes (Hyman et al., 2022). These characteristics make the York River an ideal natural laboratory for nursery habitat comparisons among multiple size classes.

The river was divided into sections based on morphology, constituting downriver, midriver, and upriver strata (Fig. 1). This is in contrast to Hyman et al. (2023), which arbitrarily divided the system into three approximately equal sections. Here, we used the Coleman Bridge (Lat = 37.2421, Long = -76.5068) as a logical delineation between downriver and midriver strata due to substantial changes in hydrology associated with this choke-point (Stockhausen and Lipcius, 2003). Delineation between midriver and upriver remained consistent with Hyman et al. (2023).

**FIG 1.**
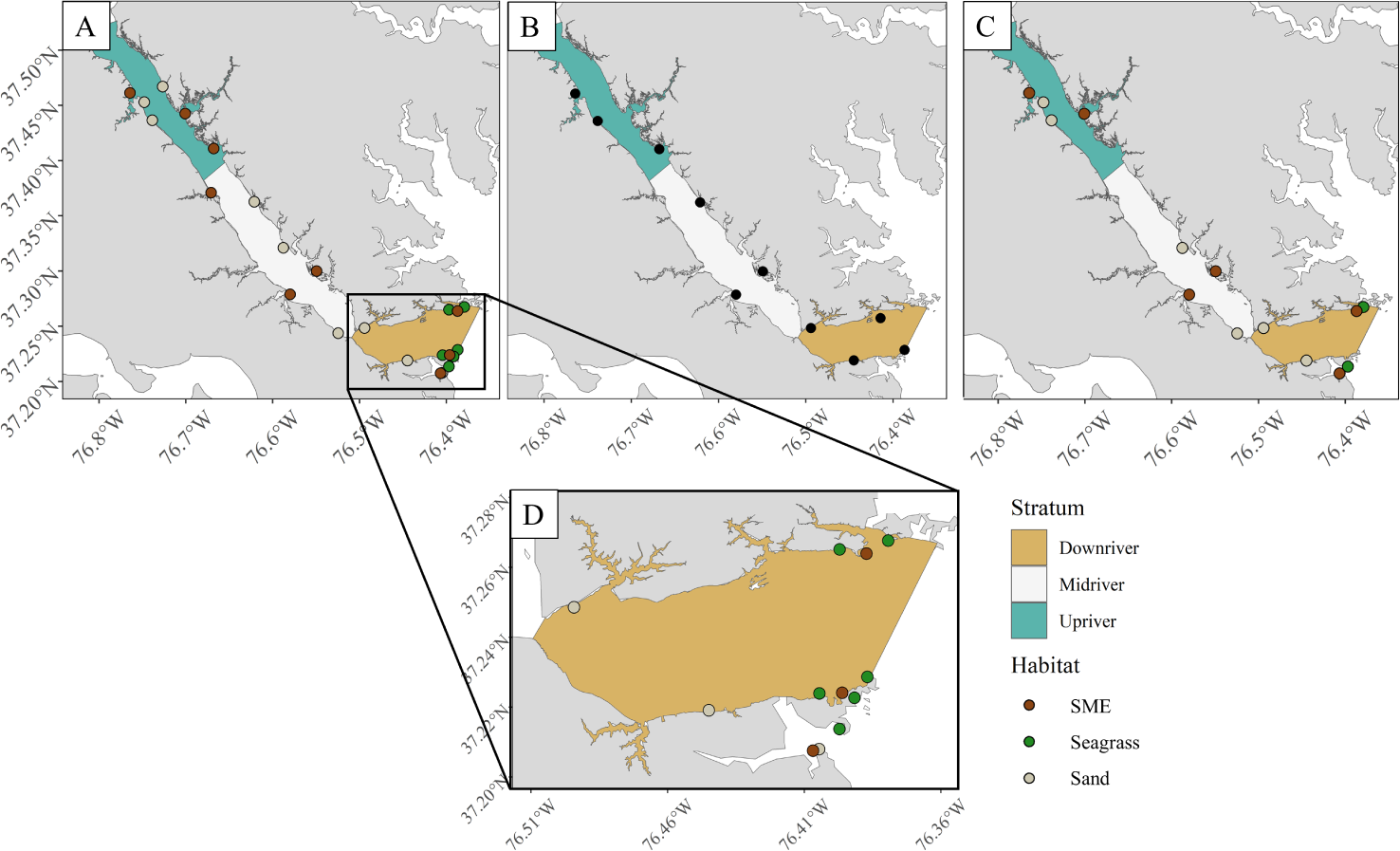
*Map displaying sampling sites for the York River. **A**: abundance sites; B: megalopae sites; **C**: survival sites; and **D**: a close-up of seagrass abundance sites sampled in the downriver stratum. Megalopae (i.e. postlarvae) sites are not habitat-specific and are not color-coded by habitat.*

### 2.2. Predictors

We considered six environmental and biological variables (herein, predictors) as potential determinants of abundance and survival for juvenile blue crabs (Tables 1 and 2). Although most predictors were included in both the abundance and survival models (Sections 2.4.1 and 2.4.2), some were only included in one model or the other. We justify the inclusion or exclusion of each predictor for a given model in Tables 1 and 2. We did not include salinity as a predictor in either model due to substantial collinearity with turbidity and location along the river axis. For a detailed description and justification of all predictors, see Appendix A.

**TABLE 1.** Descriptions and justifications of predictors used in modeling juvenile abundance.

| Predictor | Levels (if categorical) | Justification and prediction | References |
| --- | --- | --- | --- |
| Habitat | Seagrass | Abundance should be higher in structurally complex seagrass and SME relative to sand | Orth and van Montfrans (1987); Lipcius et al. (2005, 2007); Johnston and Lipcius (2012); Ralph et al. (2013); Hyman et al. (2022, 2023) |
|  | SME |  |  |
|  | Sand |  |  |
| Stratum | Downriver | Spatial position may influence abundance through spatially correlated, unobserved variables. | Posey et al. (2005); Lipcius et al. (2005) |
|  | Midriver |  |  |
|  | Upriver |  |  |
| Megalopae | continuous | Abundance is initially dictated by postlarval supply | Etherington, Eggleston and Stockhausen (2003); Heck Jr, Coen and Morgan (2001); Heck Jr and Spitzer (2001) |
| Turbidity | continuous | Abundance is positively correlated with turbidity due to higher food availability and refuge afforded by turbid habitat | Cyrus and Blaber (1987); Marley et al. (2020); O'Brien, Slade and Vinyard (1976); Howson (2000); Horodysky et al. (2010) |

**TABLE 2.**
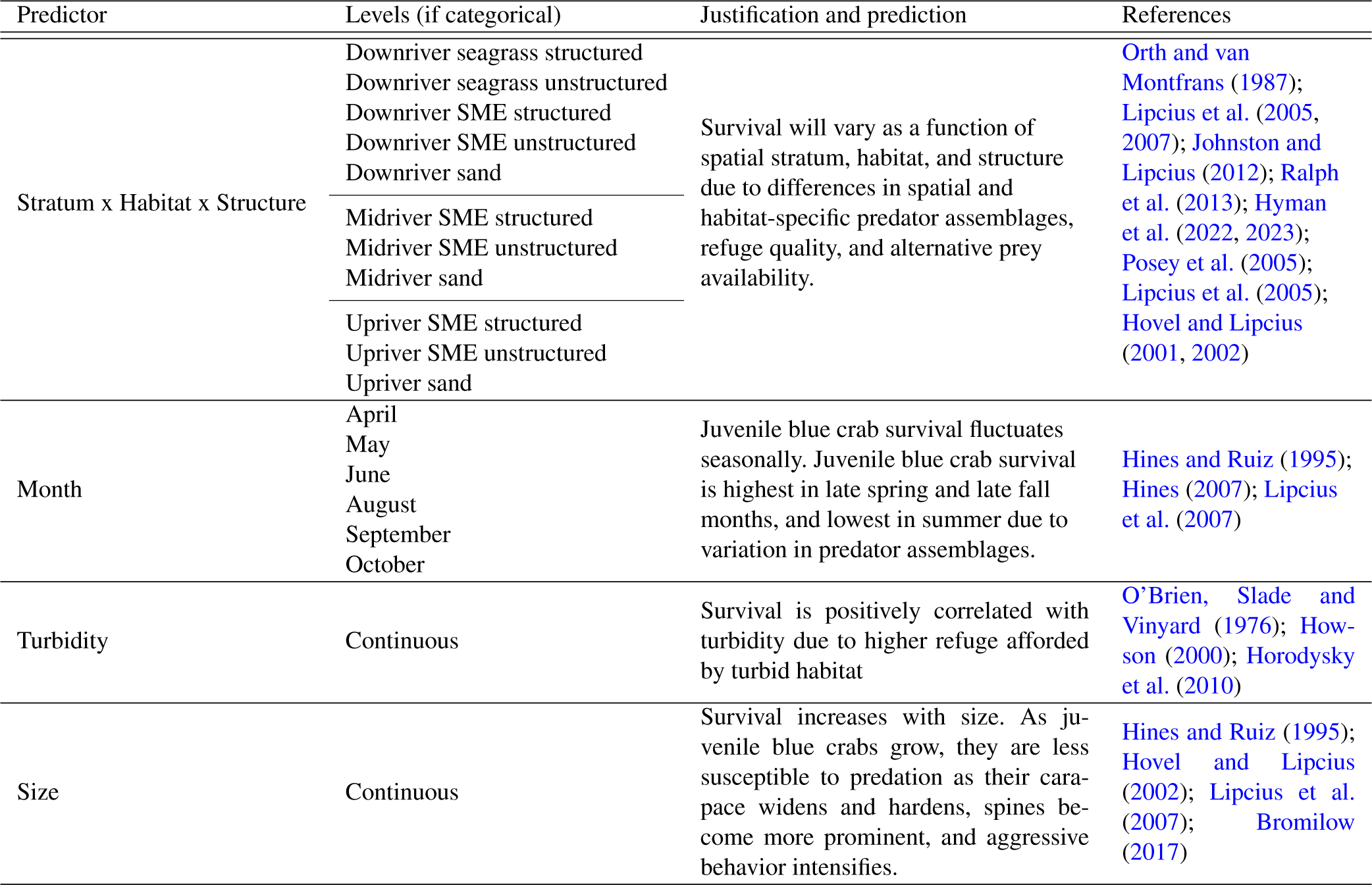
Descriptions and justifications of predictors used in the juvenile survival model. The categorical variables stratum, habitat, and structure form an incomplete, crossed design and therefore are collapsed into a single categorical variable. For details, see Appendix A.

### 2.3. Field Sampling

All sampling sites were selected from a subset of sites used in the random sampling design from Hyman et al. (2023) (Fig. 1). Subsampling was due to changes in the primary gear type for sand habitat (i.e. boat-mounted scrapes to seine hauls) which reduced the number of samples logistically feasible. Survival sites were also subsampled for logistical reasons. For juvenile abundance sampling, three SME and sand sites were randomly selected from the six habitat-specific sites in each stratum from Hyman et al. (2023) (n_sites_ = 24). The six seagrass sites used in the downriver stratum were consistent with those employed in Hyman et al. (2023). In midriver and upriver strata, seagrass was not present and only sand and SME were investigated. Megalopae sampling was used to assess the effect of postlarval supply on juvenile blue crab abundance, particularly for the small size class. Logistical limitations prevented us from sampling megalopae at every abundance site. Therefore, site selection for megalopae sampling and survival assessment was based on a random subsample of the 24 abundance sites. Specifically, three random shoreline sites in the upriver and midriver strata and four sites in the downriver stratum were randomly chosen (n_sites_ = 10). See Appendix A for a description of megalopae data preparation. Finally, in the survival study two tethering sites were randomly selected from the three abundance sites for each habitat within each stratum (n_sites_ = 14). Physicochemical variables salinity, temperature, and turbidity were recorded using a YSI data sonde (for salinity and temperature) and a Secchi disk (proxy for inverse of turbidity) at each site on each trip for all three sampling procedures. For a detailed breakdown of sampling effort by study type, habitat, and stratum, see Table A1.

#### 2.3.1. Juvenile abundance

Juvenile abundance was sampled at each station in seagrass, SME, and sand at biweekly intervals between August 5^th^ and October 14^th^, 2021 (5 trips x 24 sites = 120). Juvenile blue crabs in sand sites were sampled using a 5 m seine net (half-circle sweeps), while SME sites were sampled using modified flume nets (McIvor and Odum, 1986). At seagrass sites, a suction sampler, modified from Orth and van Montfrans (1987), was utilized to collect juvenile blue crabs (e.g., Orth and van Montfrans, 1987; Ralph et al., 2013; Hovel and Lipcius, 2002; Heck Jr, Coen and Morgan, 2001). All gear types used 3 mm^2^ mesh. Juvenile crabs were counted and measured for carapace width (see Hyman et al., 2023, for additional details on abundance sample processing).

As different sampling methods were employed for the three habitat types, gear efficiency estimates were required to scale abundance estimates for each sample. Efficiency of the suction sampling methodology is estimated at 88% (Orth and van Montfrans, 1987), while efficiency tests of the modified flume net design using marked blue crabs in fall of 2020 suggested an estimated efficiency of 92% (Hyman et al., 2023). Finally, literature suggested efficiency estimates for seine nets targeting juvenile blue crabs varied between 10–50% (mean 30%; Davis et al., 2005). These efficiency estimates and their associated uncertainties were included as Bayesian prior distributions in juvenile abundance models both to more accurately determine the effects of habitat as well as to incorporate relevant uncertainty into model estimates. For details, see Table 3, Section 2.4.1, and Hyman et al. (2023).

**Table 3.** Descriptions of predictor coefficients used in the juvenile abundance model. Prior distributions are on the model (log) scale. All β terms refer to priors of a given coefficient for all three size classes.

| Predictor | Regression Coefficient | Description | Prior (CW: $\leq 15$ mm) | Prior (CW: 16 to 30 mm) | Prior (CW: 31 to 60 mm) |
| --- | --- | --- | --- | --- | --- |
| — | $\beta_{0i}$ | Intercept of model (i.e. intercept of the reference, taken to be sand downriver) | $N(-1.10, 0.75)$ | $N(-0.11, 0.67)$ | $N(-0.11, 2.00)$ |
| Turbidity | $\beta_{1i}$ | Effect of water cloudiness measured as the negative log transformation of the Secchi disk depth (m) in site $s$ on date $t$ (see Appendix A for details) | $N(0.19, 0.19)$ | $N(0.29, 0.17)$ | $N(0.29, 2.00)$ |
| SME | $\beta_{2i}$ | Effect of SME relative to the reference | $N(2.59, 0.51)$ | $N(2.72, 0.4)$ | $N(2.72, 2.00)$ |
| Seagrass | $\beta_{3i}$ | Effect of seagrass habitat relative to the reference | $N(3.67, 0.64)$ | $N(2.07, 0.49),$ | $N(2.07, 2.00)$ |
| Midriver | $\beta_{4i}$ | Effect of midriver stratum relative to the reference | $N(0, 10)$ | $N(0, 10)$ | $N(0, 10)$ |
| Upriver | $\beta_{5i}$ | Effect of upriver stratum relative to the reference | $N(1.20, 10)$ | $N(1.2, 10)$ | $N(1.2, 10)$ |
| Megalopae | $\beta_{6i}$ | Effect of $\ln(\text{average megalopae abundance} + 1)$ in a given stratum corresponding to site $s$ four weeks prior to the $t^{\text{th}}$ date (see Appendix A for details) | $N(0, 10)$ | $N(0, 10)$ | $N(0, 10)$ |
| Seine efficiency | $\beta_{7i}$ | Suction efficiency parameter for juvenile blue crab abundance in seagrass | $N(-0.13, 0.02)$ | $N(-0.13, 0.02)$ | $N(-0.13, 0.02)$ |
| Suction efficiency | $\beta_{8i}$ | Flume efficiency parameter for juvenile blue crab abundance in SME | $N(-0.08, 0.02)$ | $N(-0.08, 0.02)$ | $N(-0.08, 0.02)$ |
| Seine efficiency | $\beta_{7i}$ | Seine efficiency parameter for juvenile blue crab abundance in sand | $N(-1.27, 0.41)$ | $N(-1.27, 0.41)$ | $N(-1.27, 0.41)$ |
| Area (log) | $A$ | Offset term relating juvenile blue crab abundance to surface area of gear used to sample habitat $h$ | — | — | — |

#### 2.3.2. Megalopae

Megalopae sampling was used to assess the effects of postlarval supply on juvenile blue crab abundance. Sampling took place at biweekly intervals between July 16^th^ and October 7^th^, 2021, following new and full moon cycles (Van Montfrans et al., 1995; Epifanio, 2007, 2019, 7 trips x 10 sites = 70). Collectors consisted of a hog’s hair filter sleeve surrounding an inner PVC cylinder (0.18 m^2^; Van Montfrans et al., 1995; Metcalf et al., 1995), deployed for approximately 12 h from sunset to sunrise. Upon collection, filters were submersed in fresh water in the field for 1 h to remove megalopae. The filters were then transported to the lab and rinsed again in fresh water three times until no fauna remained, and subsequently sieved using 500 micron mesh. Sieve contents were sorted underneath a magnifying glass for blue crab megalopae, which are distinct in coloration and morphology relative to other local estuarine crustacean larvae (Ogburn et al., 2011). Megalopae were counted and recorded along with local physicochemical variables as described in Appendix A.

#### 2.3.3. Survival

A tethering experiment was conducted at biweekly intervals between April and November, 2021 with juvenile crabs of 6–50 mm CW (n = 848), using an established tethering technique to assess survival (see Lipcius et al., 2005, for details). July was not considered because logistical issues prevented sampling in that month. Tethering was conducted in *<*1 m mean low water to limit the influence of depth (e.g. Ruiz, Hines and Posey, 1993). At each site, five crabs were haphazardly selected and tethered in both structured (where present) and unstructured treatments. Within a habitat/treatment, individual tethers were haphazardly spaced *∼*5 m apart. The size (CW) of each crab was measured to the nearest 0.1 mm using calipers prior to deployment, and deployed for *∼*24 h. Within SME and seagrass, locations within the delineated habitat which were devoid of vegetation were regarded as “unstructured” and used to compare variation in survival at the patch scale. Unstructured SME habitat was defined as areas devoid of vegetation immediately adjacent to the SME, whereas unstructured seagrass habitat was defined as interstitial barren patches within or immediately adjacent to seagrass beds. In contrast, “structured” SME and seagrass habitat were defined as localities within those habitats where vegetation was present. Within SME and seagrass habitats, crabs were tethered in both structured and unstructured treatments. Only seagrass patches with 100% aerial cover were considered in the structured seagrass treatment, while sand consisted of only an unstructured treatment. Additional details, including escape testing and assessment of treatment-specific bias (e.g. Peterson and Black, 1994; Baker and Waltham, 2020), can be found in Appendix B.

### 2.4. Analysis

All data analyses, transformations, and visualizations were completed using the R programming language for statistical computing (R Core Team, 2022a) and the Stan probabilistic programming language for Bayesian statistical modeling (Stan Development Team, 2022b, 2020).

#### 2.4.1. Abundance

Relationships between abundance and environmental variables for small juvenile blue crabs were modeled using a multivariate negative binomial linear mixed-effects model under a Bayesian framework. Predictor variables for juvenile abundance data included habitat (seagrass, SME, and sand), stratum (downriver, midriver, and upriver), megalopae local abundance four weeks prior to each sampling trip (averaged across stratum), and turbidity. Natural log (ln)-transformations were applied to both turbidity and megalopae abundance prior to analyses (see Appendix A for details).

Extending the model from Hyman et al. (2023), for the *s*^th^ site on date *t* in habitat *h*, the model for juvenile blue crab abundance in the *i^th^* size class is expressed as:

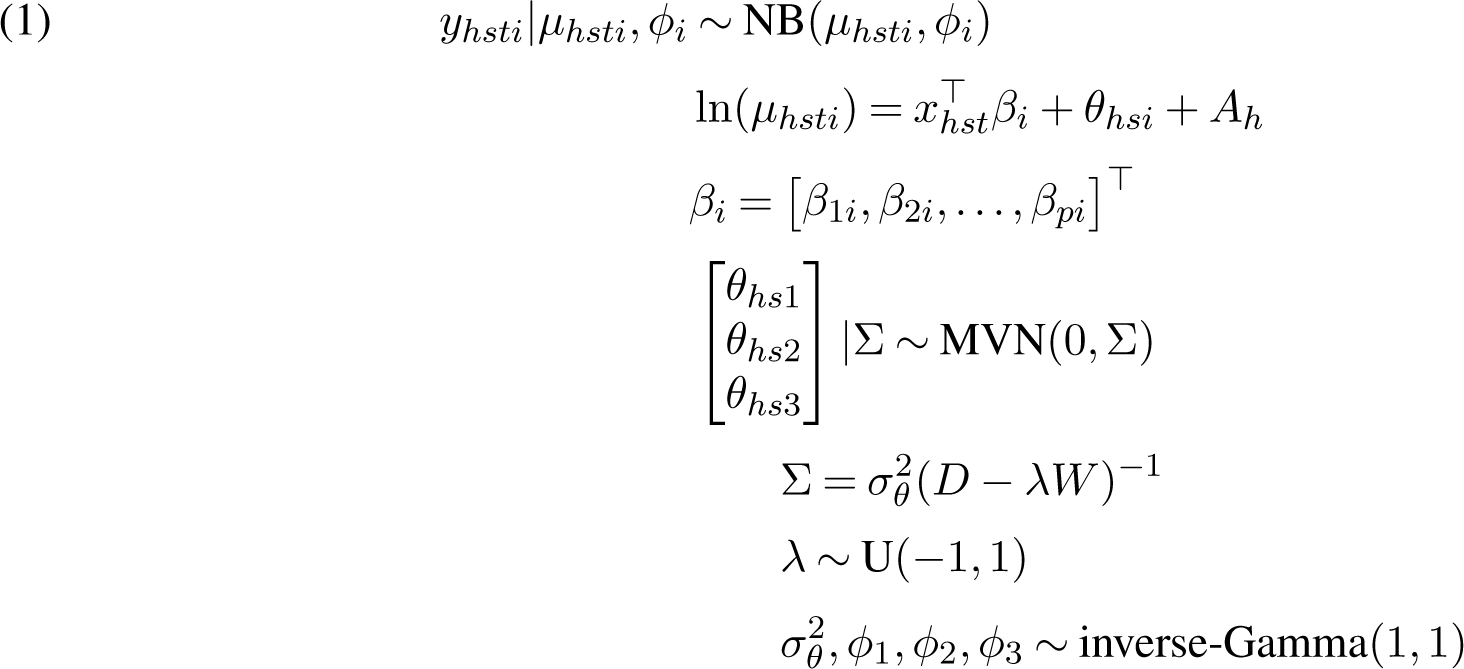

where *NB*(*µ_hsti_, ϕ_i_*) denotes a negative binomial type II distribution with mean *µ_hsti_*, while *ϕ_i_* controls the over-dispersion for each size class such that *E*[*y_hsti_*] and 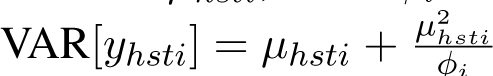. The *i* response variable, juvenile crab counts for each size class *i*, is denoted *y_hsti_* where *i* = 1 denotes CW *≤*15 mm, *i* = 2 denotes 16–30 mm, and *i* = 3 denotes 31–60 mm. Total area sampled in habitat *h* (SME = 1 m^2^, seagrass = 1.68 m^2^, sand = 9.81 m^2^) is included as an offset term *A_h_*. Meanwhile, *β_i_* refers to regression coefficients for each size class *i* associated with predictors *x_hst_*. Measurements of both the abundances of size classes and predictors *x_hst_* were taken at the site-trip spatiotemporal resolution, such that predictors were not specific to any one size class *i* but to all sizes classes at a given site-trip. Here, *θ_hsi_* denotes a site-specific random effect for a given size class *i*. The joint probability distribution of (*θ_hs_*_1_*, θ_hs_*_2_*, θ_hs_*_3_) is specified as multivariate normal with a mean vector of 0s and variance-covariance matrix Σ. The Σ matrix describes dependence among size classes based on the nearest neighbor structure specified by a 3 *×* 3 adjacency matrix, *W*, and an autocorrelation parameter *λ*, which controls the degree of autocorrelation among size classes. We employed a binary weighting scheme for *W* where *w_i,i_′* = 0 for all (*i, i^′^*) unless size classes *i ̸*= *i^′^* were adjacent. For example, the smallest size class (*≤*15 mm CW) and the next largest *i* = 2 (16–30 mm CW) are considered adjacent because increases in size among individuals in *i* = 1 would shift them to *i* = 2, whereas size classes *i* = 1 and *i* = 3 are not considered adjacent because individuals in size class *i* = 1 (*≤*15 mm CW) would need to move through size class *i* = 2 prior to reaching *i* = 3 (31–60 mm CW). Hence the 3 *×* 3 binary adjacency matrix employed here is expressed as:

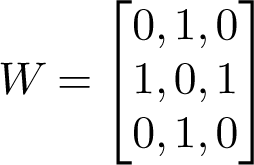

The influence of an adjacent size class on a given size class was standardized by subtracting *λW* from *D*, a diagonal matrix where *D_i,i_* is the number of neighbors for size class *i* (1, 2, and 1 for size classes *i* = 1, 2, and 3 respectively). The parameter *λ* was constrained between -1 and 1 through a uniform prior. This size class dependence structure was assumed to be homoscedastic through the variance parameter *σ*^2^, with an inverse-Gamma(1, 1) hyperprior. A similar parameterization is outlined in Hyman et al. (2022), although here the nearest neighbor structure refers to covariance among size classes instead of covariance across spatial polygons. Results from Hyman et al. (2023) indicated that spatial dependence was accounted for by spatial stratum and site random effects.

Initial model priors for fixed-effects coefficients were derived from the posterior inference from a study in the York River from 2020 examining the same sites, habitats, and size classes except for the largest size class *i* = 3 (Hyman et al., 2023). For size class *i* = 3 (31–60 mm CW), supplied prior means were identical to those for size class *i* = 2, while prior variances were scaled by a factor of 4 (model scale) to account for higher prior uncertainty associated with this size class. Descriptions of fixed effects included in the preliminary juvenile abundance model as well as their corresponding prior distributions are listed in Table 3.

#### 2.4.2. Survival

Crab survival, recorded as 1 (alive) or 0 (eaten), was analyzed for probability of survival using a hierarchical logistic regression mixed-effects model. Within stratum, unstructured treatments among habitats (seagrass, SME, and sand for downriver or SME and sand otherwise) were compared to isolate effects of differing predation pressure and refuge. Within a given stratum and within SME and seagrass habitats (where present), nearby structured and unstructured treatments were compared to facilitate inference on the effect of structure while controlling for differences in predation pressure. Due to the nature of our sampling design, we approximated nonlinear effects of seasonality by using month as a categorical fixed effect.

For the *j*^th^ tether trial in the *s*^th^ site on date *t*, the model for juvenile blue crab survival is expressed as:

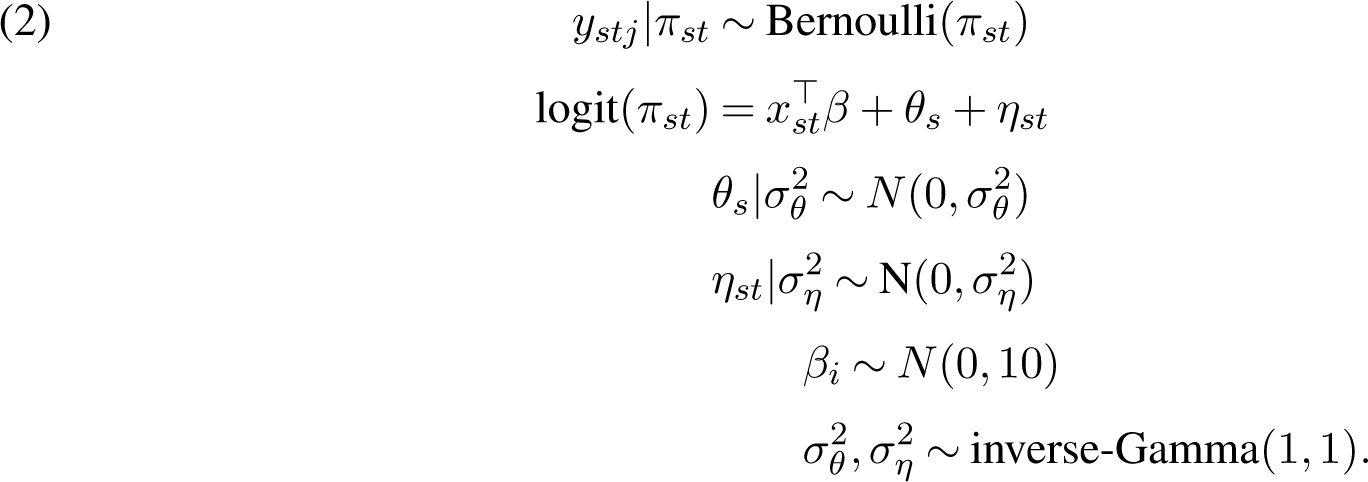

The response, binary juvenile crab survival *y_stj_* for the *j*^th^ tethering trial, is distributed as a Bernoulli random variable with probability of survival, *π_st_*. Tethering trials entailed repeatedly using the same sites, which may introduce site-specific bias. Moreover, 5 m spacing of individual tethers within a site may not have been sufficient to guarantee independence (Baker and Waltham, 2020). Hence, *θ_s_* and *η_st_* denote site-specific and site within trip-specific random effects, respectively. All regression coefficients were assigned diffuse priors (*N* (0, 10)). Descriptions of the fixed effects for the preliminary juvenile blue crab survival model can be found in Table 4.

**TABLE 4.** Descriptions of regression coefficients used in the juvenile survival model. The categorical variables stratum, habitat, and structure form an incomplete, crossed design and therefore are collapsed into a single categorical variable. For details, see Appendix A.

| Predictor | Regression Coefficient | Description |
| --- | --- | --- |
| — | $\beta_0$ | Intercept of model (i.e. the reference intercept, taken to be the intercept for sand downriver in April) |
| Turbidity | $\beta_1$ | Effect of water cloudiness measured as the negative log transformation of the Secchi disk depth (m) in site $s$ on date $t$ (see Appendix A for details) |
| Crab width | $\beta_2$ | Effect of crab width (mm) |
| May | $\beta_3$ | Effect of May relative to the reference |
| June | $\beta_4$ | Effect of June relative to the reference |
| August | $\beta_5$ | Effect of August relative to the reference |
| September | $\beta_6$ | Effect of September relative to the reference |
| October | $\beta_7$ | Effect of October relative to the reference |
| Downriver SME structured | $\beta_8$ | Effect of downriver structured SME relative to the reference |
| Midriver SME structured | $\beta_9$ | Effect of midriver structured SME relative to the reference |
| Upriver SME structured | $\beta_{10}$ | Effect of upriver structured SME relative to the reference |
| Downriver SME unstructured | $\beta_{11}$ | Effect of downriver unstructured SME relative to the reference |
| Midriver SME unstructured | $\beta_{12}$ | Effect of midriver unstructured SME relative to the reference |
| Upriver SME unstructured | $\beta_{13}$ | Effect of upriver unstructured SME relative to the reference |
| Downriver seagrass structured | $\beta_{14}$ | Effect of downriver structured seagrass relative to the reference |
| Downriver seagrass unstructured | $\beta_{15}$ | Effect of downriver unstructured seagrass relative to the reference |
| Midriver sand | $\beta_{16}$ | Effect of midriver sand relative to the reference |
| Upriver sand | $\beta_{17}$ | Effect of upriver sand relative to the reference |

### 2.5. Model implementation and validation

For each model, Bayesian inference required numerical approximation of the joint posterior distribution of all model parameters including the vectors of random effects. To this end, we implemented the above models using the Stan programming language for Bayesian inference to generate Hamiltonian Monte Carlo (HMC) samples from the posterior (Gelman, Lee and Guo, 2015). For each model, we ran four parallel Markov chains, each with 5,000 iterations for the warm-up/adaptive phase, and another 5,000 iterations as posterior samples (i.e. 20,000 draws in total for posterior inference). Convergence of the chains was determined both by visual inspection of trace plots (e.g. Fig. A1) and through inspection of the split *R*^^^ statistic. All sampled parameters had an *R*^^^ value less than 1.01, suggesting chain convergence (Gelman, Lee and Guo, 2015). Covariates and interactions whose regression coefficients had Bayesian confidence intervals (CIs) that excluded 0 at a confidence level of 80% were considered scientifically relevant to juvenile blue crab abundance and survival. All CIs referenced here are highest posterior density intervals (McElreath, 2018).

### 2.6. Conditional inference

Conditional means and conditional effects plots were used to assess the relationship between response variables (juvenile blue crab abundance and survival) and meaningful predictors both among habitats within a size class and within habitats between size classes. Herein, we refer to “conditional” as holding all random effects at 0 and fixing co-varying predictors. For a detailed description of the estimation procedure for each conditional quantity, see Table 5. Conditional effects for categorical terms are reported using posterior median values and 80% CIs, while the relationship between the conditional mean (*µ*_cond_*_vi_* or *π*_cond_*_v_*) and each continuous predictor (*x_vi_* or *x_v_*) was plotted with posterior medians as well as 50%, 60%, and 80% credible bands.

**TABLE 5.** Descriptions of conditional means and conditional effects derived from the abundance and survival models.

| Model | Term | Description | Equation |
| --- | --- | --- | --- |
| Abundance | $\mu_{\text{cond}_{hi}}$ | The expected number of crabs in size class $i$ in habitat $h$, with random effects fixed at 0, ln turbidity and ln megalopae fixed at 0, and stratum taken to be downriver (reference). | $\exp(\beta_{0i})$ for $h = \text{reference}$<br>$\exp(\beta_{hi} + \beta_{0i})$ for $h > \text{reference}$ |
| | $\mu_{\text{cond}_{vi}}(x_{vi})$ | The expected number of crabs in size class $i$ as a function of continuous predictor $x_{vi}$, with random effects fixed at 0, all other continuous predictors fixed at 0, and categorical terms fixed at the reference (i.e. downriver sand). | $\exp(\beta_{0i} + \beta_{vi}x_{vi})$ |
| | $\eta_{\text{cond}_{hi}}$ | The $\ln \mu_{\text{cond}_{hi}}$ (i.e. the linear predictor). | $\beta_{0i}$ for $h = \text{reference}$<br>$\beta_{hi} + \beta_{0i}$ for $h > \text{reference}$ |
| | $L_{hi-mi}$ | The linear contrast between habitat $h$ and habitat $m$ for size class $i$ | $\eta_{\text{cond}_{hi}} - \eta_{\text{cond}_{mi}}$ |
| Survival | $\pi_{\text{cond}_j}$ | The probability of survival at categorical level $j$, with random effects fixed at 0, ln turbidity fixed at 0, and crab width fixed at 22.4 mm (i.e. average crab size in study). When $j$ denotes a stratum-habitat-structure combination, month is held at April. When $j$ denotes a month, stratum-habitat-structure is fixed at downriver sand. | $\text{expit}(\beta_0)$ for $j = \text{reference}$<br>$\text{expit}(\beta_j + \beta_0)$ for $j > \text{reference}$ |
| | $\gamma_{\text{cond}_j}$ | The linear predictor ($\text{logit } \pi_{\text{cond}_j}$) | $\beta_0$ for $j = \text{reference}$<br>$\beta_j + \beta_0$ for $j > \text{reference}$ |
| | $W_{j-r}$ | The linear contrast between levels $j$ and $r$. | $\gamma_{\text{cond}_j} - \gamma_{\text{cond}_r}$ |
| | $\pi_{\text{cond}_v}(x_v)$ | The probability of survival as a function of continuous predictor $x_v$, with random effects fixed at 0, all other continuous predictors fixed at 0 (except crab CW, fixed at 1 mm to show increased contrast in survival probabilities), and categorical terms fixed at the reference. | $\text{expit}(\beta_0 + \beta_v x_v)$ |

## 3. Results

### 3.1. Abundance

#### 3.1.1. Patterns in abundance among small-sized (≤15 mm) juveniles

Habitat was an important driver of small juvenile blue crab abundance (Table 6; Fig. A2). Among habitats, the estimate (i.e. posterior median) of the conditional mean (*µ*_cond_*_h,_*_15_, Table 5) for small juvenile blue crab abundance was highest in seagrass, followed by SME and sand (Table 6; Fig. 2, left panel). Linear contrasts (*L_h,_*_15_*_−m,_*_15_) between habitats for small juvenile blue crabs all yielded 80% Bayesian CIs that excluded 0, indicating that differences in the expected number of small juvenile crabs among habitats were statistically meaningful (Fig. 3, left panel). Herein, we use the term “indicate” to refer to inferences with strong statistical support (see section 2.5).

**TABLE 6.** *Posterior summary statistics (median and 80% CIs) for the small (≤15 mm CW) juvenile size class. Values under the habitat columns refer to η_condh,_*_15_ *(model scale) and µ_condh,_*_15_ *(count scale) and should be interpreted as the expected small juvenile abundance in a given habitat at a given site with 0* ln *turbidity and 0* ln *megalopae. Values under the* ln *turbidity and* ln *megalopae columns reflect abundance model slope terms (β) for those continuous predictors with categorical terms held at the reference (i.e. downriver sand). Stratum effects were not statistically meaningful for any size class and are not reported here*.

| Scale | Quantile | Sand | Seagrass | SME | $\ln$ Turbidity | $\ln$ Megalopae |
| --- | --- | --- | --- | --- | --- | --- |
| Model | 10% | -1.75 | 1.84 | 0.70 | 0.06 | 0.01 |
|  | 50% | -1.31 | 2.20 | 1.10 | 0.27 | 0.12 |
|  | 90% | -0.84 | 2.58 | 1.51 | 0.47 | 0.24 |
| Count | 10% | 0.17 | 6.27 | 2.01 | 1.06 | 1.01 |
|  | 50% | 0.27 | 9.06 | 3.02 | 1.31 | 1.13 |
|  | 90% | 0.43 | 13.16 | 4.52 | 1.61 | 1.27 |

**Fig 2.**
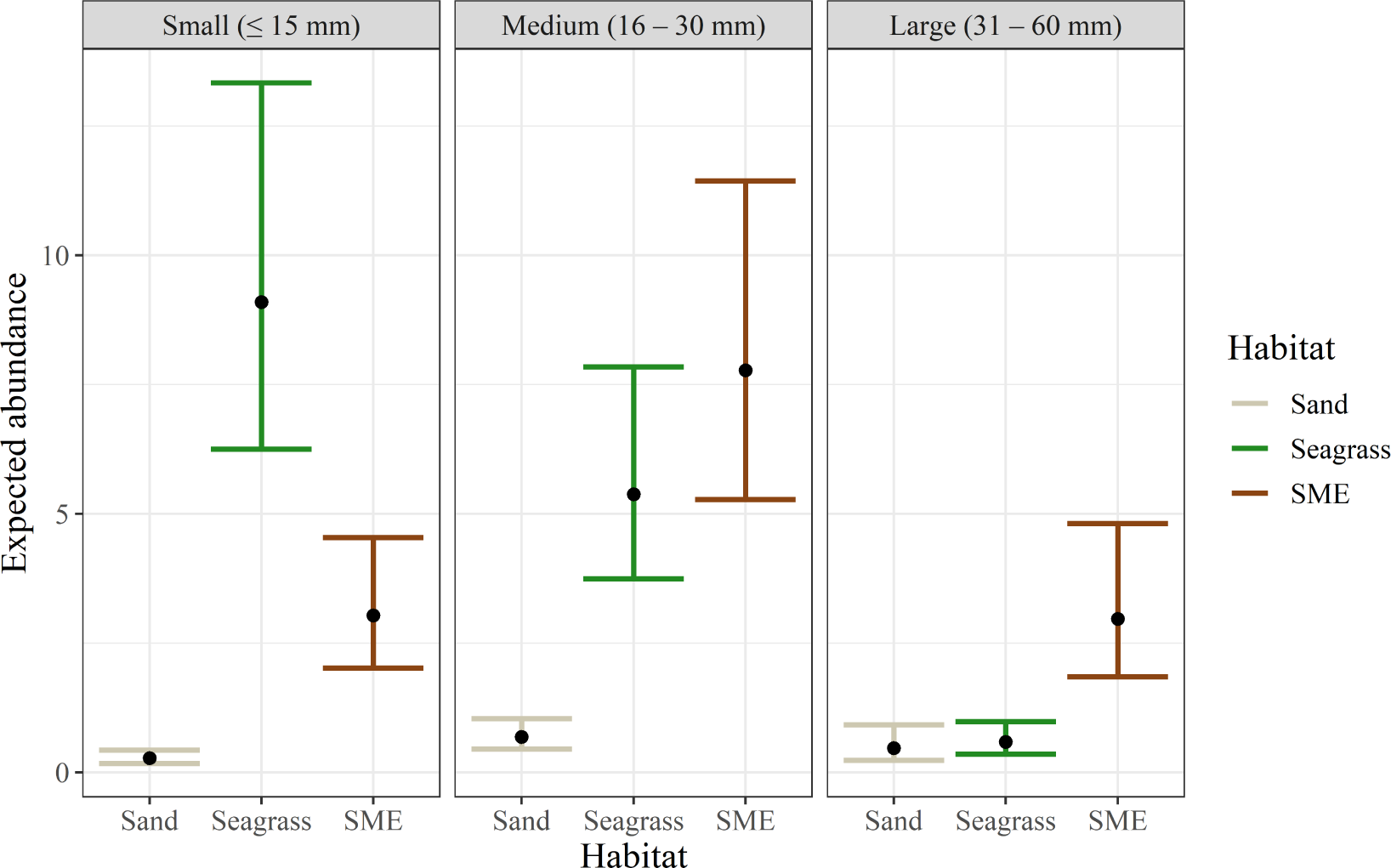
*Posterior median and 80% CI for µ_condhi_ of habitat for small (≤15 mm CW; left column), medium (16–30 mm CW; middle column), and large (31–60 mm CW; right column) size classes*.

**Fig 3.**
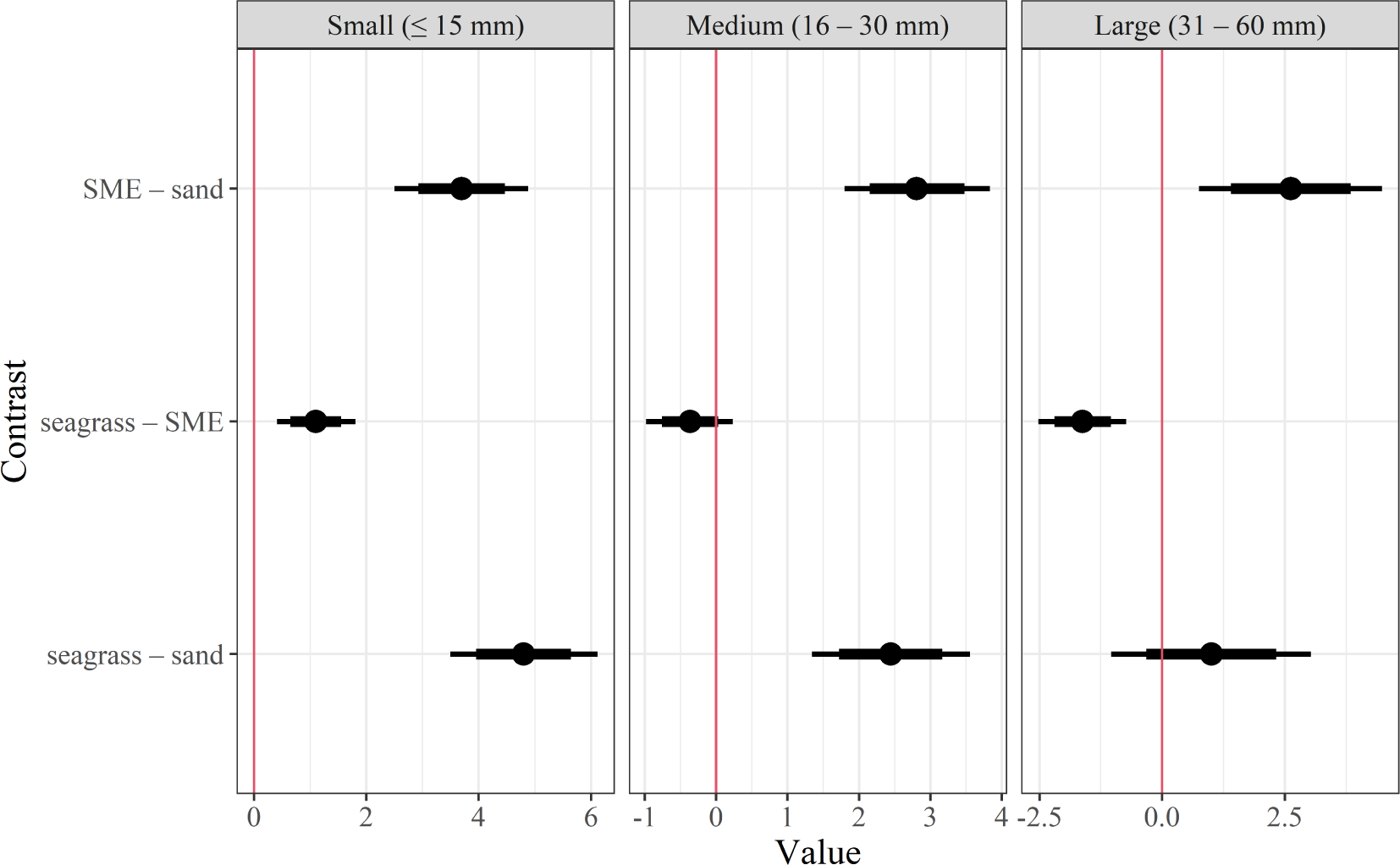
*Linear contrast statements (L_hi−mi_) depicting conditional differences in expected juvenile blue crab abundance between habitats by size class. Dots denote posterior median difference in expected values, while thick bars represent 80% Bayesian CIs and thin bars denote 95% Bayesian CIs. The red vertical line denotes 0*.

Turbidity and megalopae abundance were both positively associated with small juvenile blue crab abundance – 80% CIs of both (model scale) are above 0 (Table 6 and Fig. A2). Conversely, effects of midriver and upriver strata relative to the downriver stratum (reference) were not informative (Fig. A2). Expected small juvenile abundance increased with both average megalopae per collector (back-transformed from *ln* (megalopae +1) and ln turbidity (Figs. 4A and 4B).

**FIG 4.**
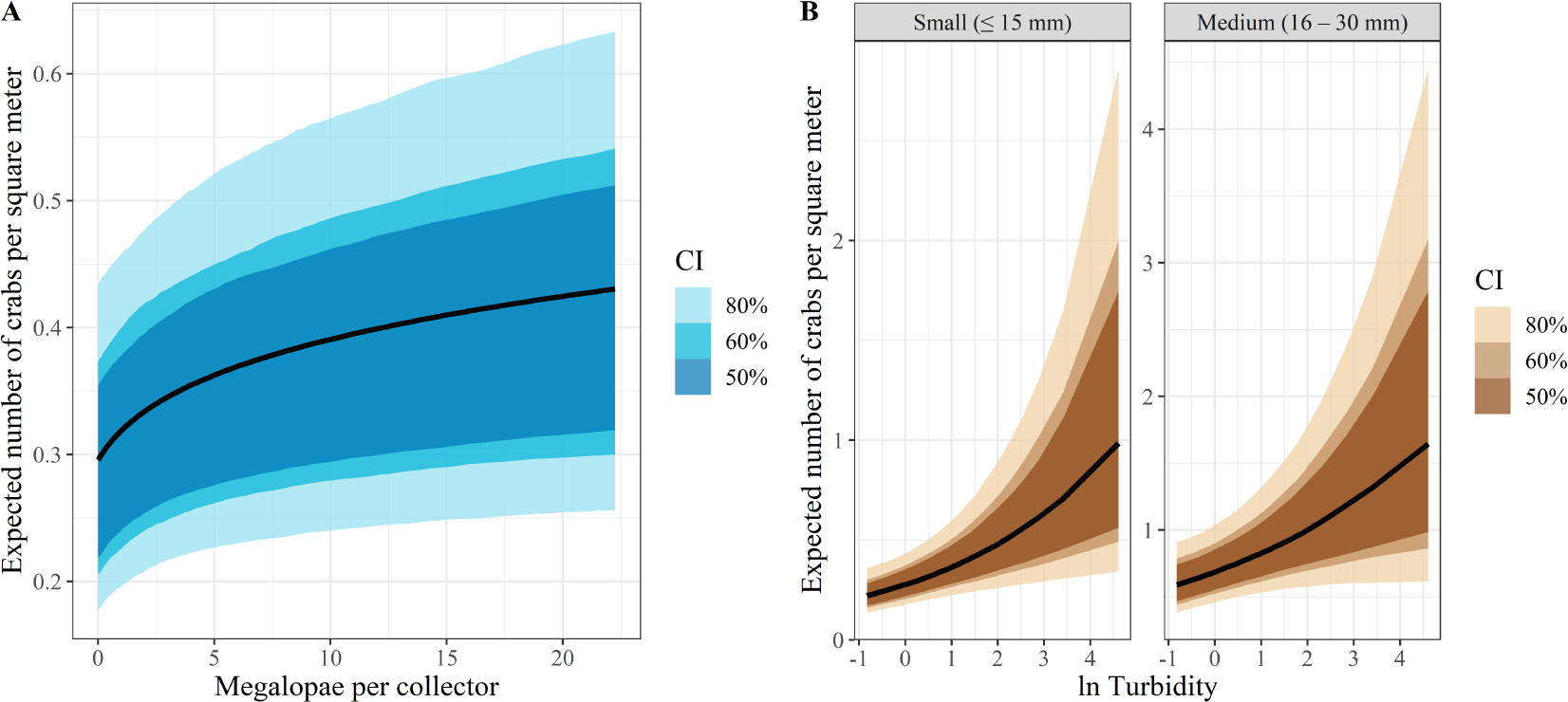
*Conditional relationships between expected juvenile blue crab abundance (µcond_vi_* (*xv*)*) as a function of **A**) average megalopae abundance per collector and **B**)* ln *turbidity. The response value for **A**) is the expected number for small size class, while the response values for **B**) include both small (left) and medium (right) size classes. Colored bands indicate credible bands ranging from 50% (0.5) to 80% (0.8) credibility*.

#### 3.1.2. Patterns in abundance among medium-sized (16–30 mm) juveniles

Habitat and turbidity were both relevant drivers of abundances of medium-sized juvenile blue crabs (Fig. A3). Based on the posterior median of *µ*_cond_*_hi_* (Table 5), unlike the smaller size class, medium-sized juveniles were most abundant in SME, followed by seagrass and sand (Table 7; Fig. 2). Similar to small juveniles, linear contrasts (*L_h,_*_30_*_−m,_*_30_) among habitats for juveniles in the medium size class indicated that differences in the expected number of medium-sized juvenile crabs among habitats were statistically meaningful, although the linear contrast between seagrass and SME was marginally meaningful (i.e. 80% CI = -0.76 to 0.031; Fig. 3, middle panel). Turbidity was also positively associated with medium-sized juvenile abundance (Table 7 and Fig. A3). However, posterior distributions of coefficients for both megalopae and spatial strata indicated that these predictors were not meaningful in predicting abundances of medium-sized juveniles (Fig. A3).

**TABLE 7.** *Posterior summary statistics (median and 80% CIs) for the medium (16–30 mm CW) juvenile size class. Values under the habitat columns refer to η_condh,_*_30_ *(model scale) and µ_condh,_*_30_ *(count scale) and should be interpreted as the expected small juvenile abundance in a given habitat at a given site with 0* ln *turbidity and 0* ln *megalopae. Values under the* ln *turbidity and* ln *megalopae columns reflect abundance model slope terms (β) for those continuous predictors with categorical terms held at the reference (i.e. downriver sand). Stratum effects were not statistically meaningful for any size class and are not reported here*.

| Scale | Quantile | Sand | Seagrass | SME | $\ln$ Turbidity | $\ln$ Megalopae |
| --- | --- | --- | --- | --- | --- | --- |
| Model | 10% | -0.79 | 1.32 | 1.67 | 0.00 | -0.15 |
|  | 50% | -0.38 | 1.68 | 2.05 | 0.19 | -0.03 |
|  | 90% | 0.05 | 2.05 | 2.43 | 0.38 | 0.11 |
| Count | 10% | 0.46 | 3.73 | 5.32 | 1.00 | 0.86 |
|  | 50% | 0.68 | 5.36 | 7.74 | 1.21 | 0.97 |
|  | 90% | 1.05 | 7.75 | 11.40 | 1.46 | 1.11 |

#### 3.1.3. Patterns in abundance of large (31–60 mm) size class

Habitat was the only predictor that was statistically informative of abundances among the largest size class of juvenile blue crabs (Fig. A4). The posterior median of large juvenile conditional abundance (*µ*_cond_*_h,_*_60_) was highest in SME habitat, and lower in sand and seagrass (Table 8; Fig. 2). The 80% CIs for linear contrasts between both SME and seagrass as well as SME and sand excluded 0, although that between seagrass and sand included 0 (Fig. 3). This indicated that SME was associated with higher abundances of large juveniles relative to seagrass and sand, which harbored equivalent abundances of large juveniles. Posterior distributions for coefficients of turbidity, megalopae, and strata indicated these variables did not influence large juvenile blue crab abundance (Fig. A4).

**TABLE 8.** Posterior summary statistics (median and 80% CIs) for the large (31–60 mm CW) juvenile size class. Values under the habitat columns refer to η_condh,60_ (model scale) and µ_condh,60_ (count scale) and should be interpreted as the expected small juvenile abundance in a given habitat at a given site with 0 ln turbidity and 0 ln megalopae. Values under the ln turbidity and ln megalopae columns reflect abundance model slope terms (β) for those continuous predictors with categorical terms held at the reference (i.e. downriver sand). Stratum effects were not statistically meaningful for any size class and are not reported here.

| Scale | Quantile | Sand | Seagrass | SME | ln Turbidity | ln Megalopae |
| --- | --- | --- | --- | --- | --- | --- |
| Model | 10% | -1.41 | -1.03 | 0.62 | -0.71 | -0.23 |
|  | 50% | -0.75 | -0.53 | 1.10 | -0.23 | -0.08 |
|  | 90% | -0.06 | -0.05 | 1.58 | 0.25 | 0.07 |
| Count | 10% | 0.24 | 0.36 | 1.87 | 0.49 | 0.80 |
|  | 50% | 0.47 | 0.59 | 3.01 | 0.80 | 0.93 |
|  | 90% | 0.94 | 0.96 | 4.84 | 1.28 | 1.07 |

### 3.2. Survival

Stratum-habitat-structure, turbidity, month, and carapace width were relevant drivers of juvenile blue crab survival (Table 9 and Fig. A5). Among stratum-habitat-structure combinations, the posterior median of the conditional probability of juvenile blue crab survival (*π*_cond_*_j_*, Table 5) was highest in downriver seagrass structured habitat, followed by midriver and upriver SME structured habitat (Fig. 5). The stratum-habitat-structure combination with the lowest posterior median conditional probability of survival was downriver seagrass unstructured habitat.

**TABLE 9.**
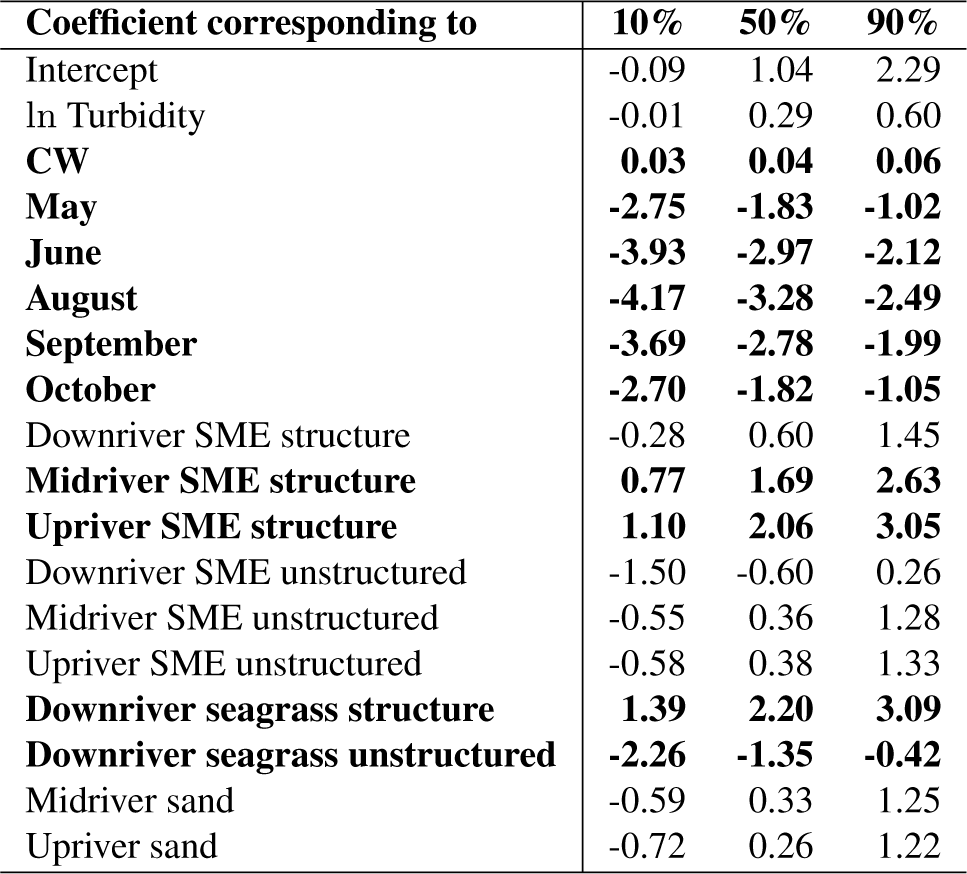
*Posterior summary statistics (median and 80% CIs) of regression coefficients β from the survival model. Effects of categorical predictors should be interpreted as relative to the reference (downriver sand in April). See* Table 4 *for descriptions of predictors. Bolded values indicate that the coefficient of a given parameter is statistically meaningful*.

| Coefficient corresponding to | 10% | 50% | 90% |
| --- | --- | --- | --- |
| Intercept | -0.09 | 1.04 | 2.29 |
| ln Turbidity | -0.01 | 0.29 | 0.60 |
| **CW** | **0.03** | **0.04** | **0.06** |
| **May** | **-2.75** | **-1.83** | **-1.02** |
| **June** | **-3.93** | **-2.97** | **-2.12** |
| **August** | **-4.17** | **-3.28** | **-2.49** |
| **September** | **-3.69** | **-2.78** | **-1.99** |
| **October** | **-2.70** | **-1.82** | **-1.05** |
| Downriver SME structure | -0.28 | 0.60 | 1.45 |
| **Midriver SME structure** | **0.77** | **1.69** | **2.63** |
| **Upriver SME structure** | **1.10** | **2.06** | **3.05** |
| Downriver SME unstructured | -1.50 | -0.60 | 0.26 |
| Midriver SME unstructured | -0.55 | 0.36 | 1.28 |
| Upriver SME unstructured | -0.58 | 0.38 | 1.33 |
| **Downriver seagrass structure** | **1.39** | **2.20** | **3.09** |
| **Downriver seagrass unstructured** | **-2.26** | **-1.35** | **-0.42** |
| Midriver sand | -0.59 | 0.33 | 1.25 |
| Upriver sand | -0.72 | 0.26 | 1.22 |

**FIG 5.**
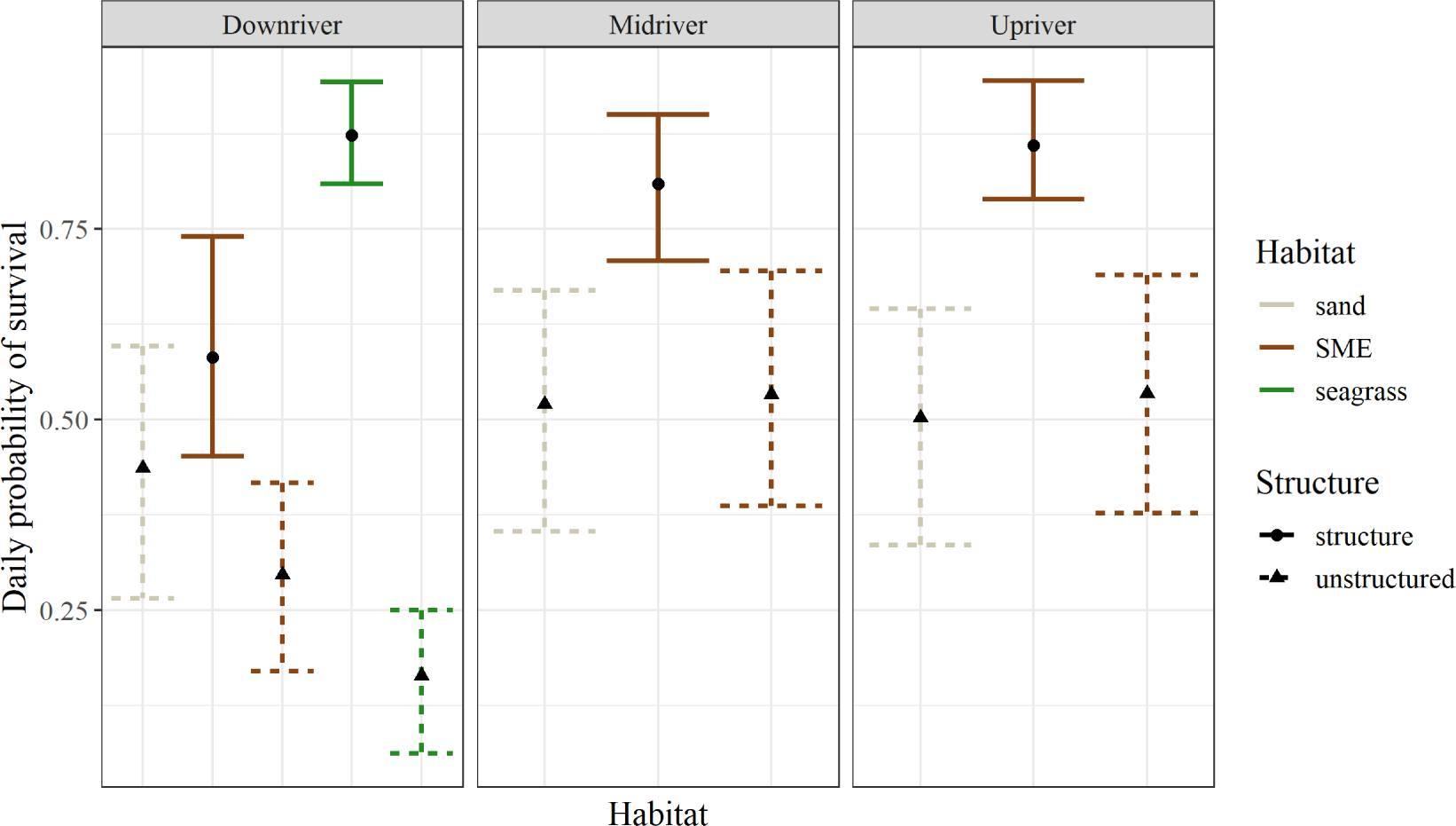
*Posterior median and 80% CIs of conditional mean (π_condj_) for habitat-structure combinations by river stratum*.

Bayesian 80% CIs of linear contrasts (*W_j−r_*) indicated that downriver structured seagrass, midriver structured SME, and upriver structured SME conferred the highest relative survival to juvenile blue crabs. Linear contrasts between these stratum-habitat-structure levels against all others were positive and excluded 0 (Table A2). However, linear contrasts among these three stratum-habitat-structure combinations indicated that they conferred equivalent probabilities of survival. Moreover, contrasts among sand habitats across all strata indicated that these habitats also conferred equivalent survival. The posterior median for the conditional probability of survival in SME habitat increased from downriver to midriver and upriver habitats. The 80% CIs of linear contrasts between downriver unstructured seagrass and all other habitats were negative and excluded 0 with the exception of downriver unstructured SME, which indicated that this habitat conferred the lowest survival.

Among months considered, the posterior median for the conditional probability of survival (*π*_cond_*_j_*) was highest in April and lowest in August (Fig. 6). Across months, this probability exhibited a nonlinear pattern in which survival peaked in spring, declined in summer months, and increased again through fall (Fig. 6). The 80% CIs of linear contrasts between April and all other months were positive and excluded 0, indicating that April conferred the highest survival among months (Fig. A6). Linear contrasts among June–September and June–August indicated that these months conferred relatively equivalent survival, although juvenile survival was lower in August than in September (Table A2 and Fig. 6). Finally, May and October conferred statistically equivalent survival which was higher than in summer months (June, August, and September) but lower than in April (Table A2 and Fig. 6).

**FIG 6.**
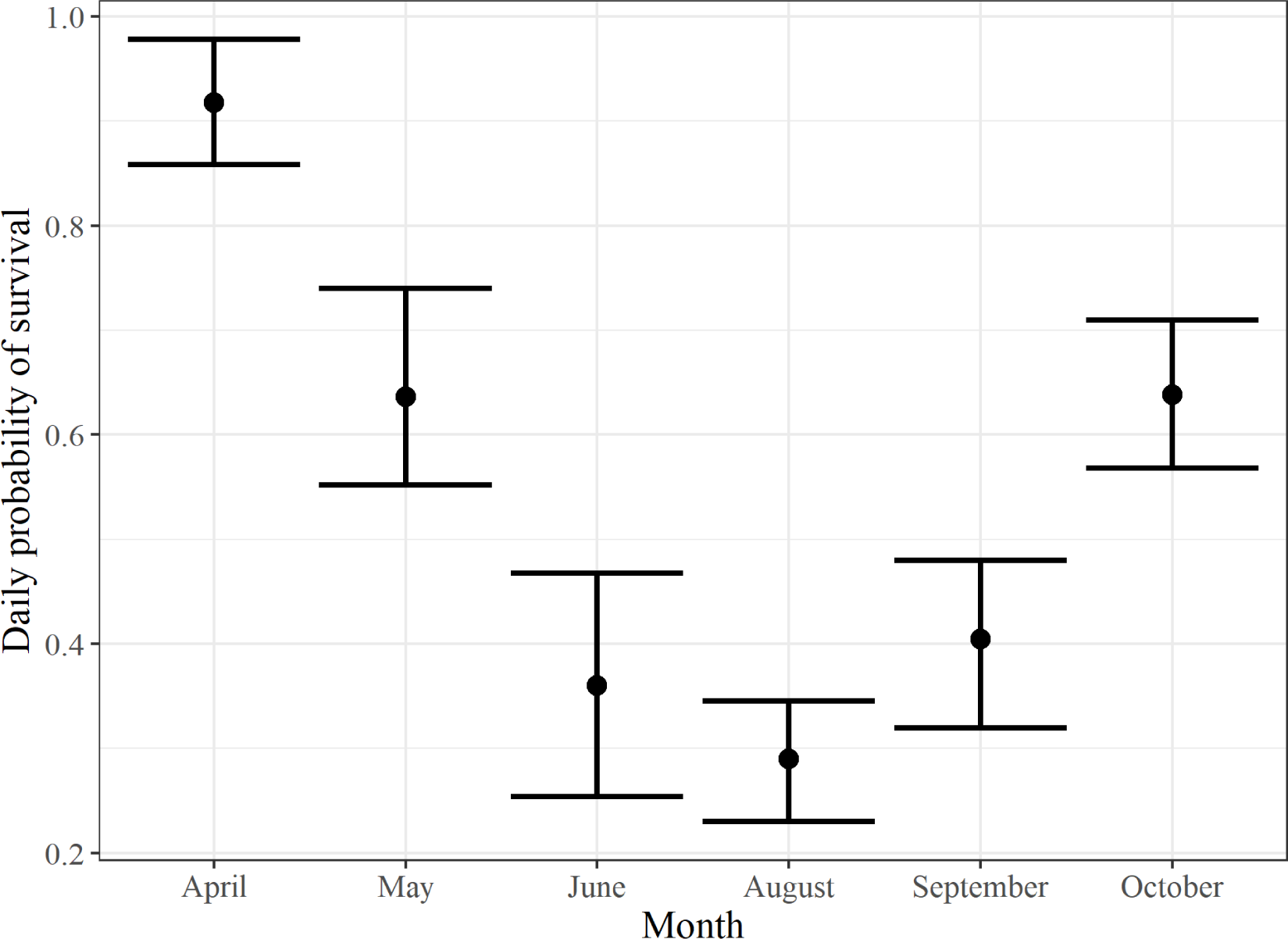
*Posterior median and 80% CIs of juvenile blue crab conditional survival probability (π_condj_) in months April, May, June, August, September, and October*.

With respect to continuous predictors, the conditional probability of survival (*π*_cond_*_v_* (*x_v_*)) increased with both carapace width and ln turbidity: the 80% CI for carapace width entirely excluded 0, which indicated CW was statistically important in explaining variation in juvenile blue crab survival. Similarly, the lower limit of the 80% CI for the effect of ln turbidity marginally contained 0 (Table 9). Based on the posterior median, the conditional probability of survival increased with ln turbidity, such that for every unit (ln(1 cm)) increase in turbidity, the odds (*<u>^π^</u>*) of survival increased by 133% (Fig. 7). Similarly, based on the posterior median, a 1-unit increase in carapace width (1 mm) corresponded with an increase in the odds of survival by 104% (Fig. 7).

**FIG 7.**
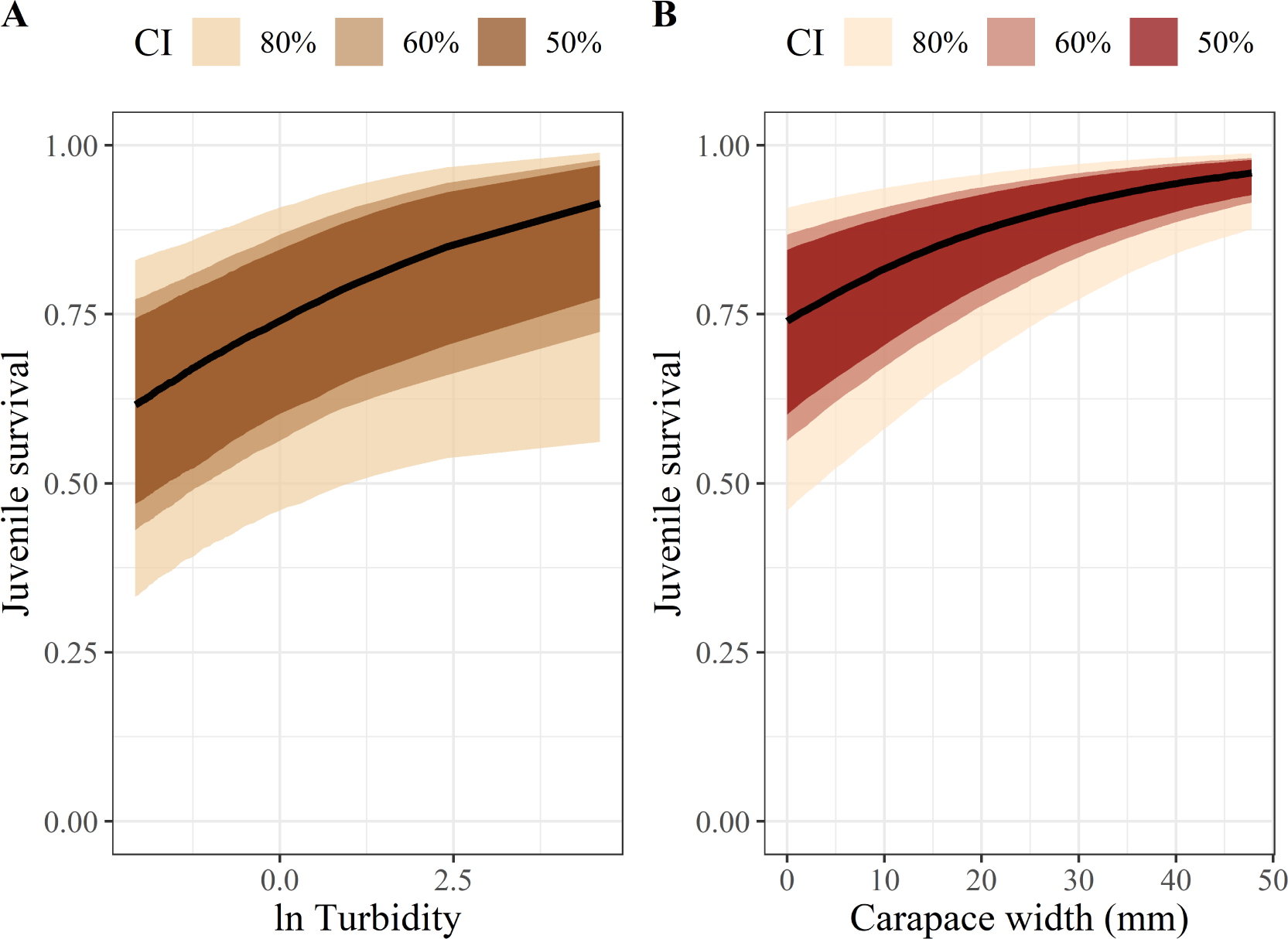
Relationships between the conditional probability of juvenile blue crab survival (πcond_v_ (xv)). Colored regions indicate credible bands.

## 4. Discussion

For a given life stage, habitat-specific abundance and survival are key determinants of a habitat’s relative importance within the seascape. Identifying and understanding the biotic and abiotic mechanisms governing these two processes both within and among habitats are critical for accurate estimation of size- and habitat-specific production. In nursery habitats of marine species, evaluation of survival, growth, abundance and ontogenetic habitat shifts has typically focused on relatively broad size ranges through the juvenile phase. Yet, ontogenetic shifts in habitat use may occur within much narrower size ranges, which has not been well studied and which is critical to the conservation and restoration of nursery habitats. Using manipulative and mensurative field experiments, we jointly assessed habitat-specific abundance and survival for multiple size classes of newly recruited juvenile blue crabs to highlight the relative importance of primary and secondary nursery habitats through ontogeny. We found that habitat-specific utilization rates differed by juvenile size class over a surprisingly narrow range of size, and were related to (1) the structural and biological characteristics of the nominal nursery habitats, (2) spatial gradients of environmental variables within the tributary, and (3) the likely trade-offs between growth and survival through ontogeny.

### 4.1. Habitat-utilization patterns of juvenile blue crabs

#### 4.1.1. Small juveniles

Habitat utilization by small (*≤*15 mm CW) juveniles was a function of habitat, postlarval supply, turbidity, and survival. Small juvenile abundance was positively correlated with megalopal recruitment, whereby small juvenile abundance sharply increased when megalopal abundance was elevated in the previous month and then tapered to an asymptote at the highest densities of megalopae. This phenomenon is consistent with density-dependent processes occurring at this size class. Strong responses to megalopal supply are expected when juvenile densities are low. However, high abundances of megalopae may exceed the carrying capacity of structurally complex nursery habitats (Heck Jr and Spitzer, 2001). As a result, post-settlement processes such as competition, cannibalism, and density-dependent secondary dispersal increasingly influence small juveniles at higher megalopal abundances (Heck Jr, Coen and Morgan, 2001; Spitzer, Heck and Valentine, 2003; Etherington and Eggleston, 2000; Reyns and Eggleston, 2004; Blackmon and Eggleston, 2001)

After accounting for variation in abundances due to megalopal supply, small juveniles were most abundant in seagrass, followed by SME and sand. High abundance of small juveniles in seagrass and SME is consistent with previous research demonstrating that structured habitats are preferred primary nurseries(seagrass; Orth and van Montfrans, 1987; Perkins-Visser, Wolcott and Wolcott, 1996; Hovel and Lipcius, 2002; Ralph et al., 2013; Johnston and Lipcius, 2012; Voigt and Eggleston, 2023) as well as alternative structured habitats (SME and complex algae; Orth and van Montfrans, 1987; Posey et al., 2005; Johnson and Eggleston, 2010; Wood and Lipcius, 2022; Hyman et al., 2022, 2023) for this life stage of juveniles.

High abundances of small juveniles in structured habitats should be considered within the context of megalopal supply. Megalopae re-enter estuaries from the continental shelf, and therefore the highest concentrations of megalopae occur near the York River mouth and decline farther upriver (Stockhausen and Lipcius, 2003). Seagrass beds only occur in the downriver portion of the York River and are positioned farther from the shoreline (and thus closer to ingressing megalopae) relative to SME. Seagrass beds are therefore most likely the first structurally complex habitat encountered by immigrating megalopae within the seascape and act as an initial settlement habitat (Stockhausen and Lipcius, 2003). Moreover, structured seagrass habitat provided the highest relative survival among all habitats considered. Survival also increased with size across habitats, adding further support that the smallest juveniles are most vulnerable to predation pressure (Pile et al., 1996; Perkins-Visser, Wolcott and Wolcott, 1996). Hence, the spatial orientation of seagrass beds relative to ingressing megalopae, coupled with the survival requirements of small juveniles, renders seagrass beds an adaptive initial settlement habitat.

SME also harbored higher abundances of small juveniles compared to sand habitat, but lower than seagrass habitat. This observation is likely a function of (1) the survival conferred by SME, (2) habitat-specific megalopal encounter probabilities, and (3) density-dependent trade-offs. First, structured SME – particularly structured SME positioned upriver – conferred roughly equivalent survival to juveniles relative to seagrass. In contrast, survival in unstructured SME was lower and roughly equivalent to sand. As access to salt marsh vegetation for aquatic organisms is controlled by marsh flooding and tidal regimes, the importance of marsh structural complexity in governing survival is likely regulated by hydrology (Minello, Rozas and Baker, 2012). In Chesapeake Bay, SME utilization is limited by mesotidal inundation profiles characteristic of mid-Atlantic estuaries (Minello, Rozas and Baker, 2012; de la Barra et al., 2022). Aggregate survival in SME is therefore a combination of the survival conferred by both structured and unstructured components of salt marshes. Indeed, the effectiveness of the flume nets used to capture juveniles in SME relied upon movement of small juveniles from structured salt marsh vegetation to unstructured bottom to remain inundated. Second, although a majority of megalopae encounter seagrass beds, a portion of megalopae miss this habitat and are advected upriver (Stockhausen and Lipcius, 2003). Here, higher survival conferred by upriver SME relative to nearby sand habitat makes SME an adaptive alternative in the absence of seagrass habitat. Finally, in downriver habitat immediately adjacent to seagrass beds, high juvenile abundance in SME habitat is likely related to density-dependent secondary dispersal of juveniles avoiding competition in seagrass beds (Etherington and Eggleston, 2000; Etherington, Eggleston and Stockhausen, 2003; Reyns and Eggleston, 2004; Blackmon and Eggleston, 2001).

#### 4.1.2. Medium-sized juveniles

In contrast to small juveniles, medium-sized (16-30 mm) juveniles were not correlated with megalopal abundance and were more abundant in SME than in seagrass, although abundances remained high in both types of structured habitats. Megalopal abundance was not expected to directly affect medium-sized or large juvenile abundances, as megalopae must transit the small juvenile stage before reaching larger size classes. Preferences of medium-sized juveniles for SME are consistent with recent findings in this system (Ralph, 2014; Hyman et al., 2023) and likely reflect shifting energetic requirements relative to predation pressure (Werner and Gilliam, 1984). In contrast to smaller juveniles, medium-sized juveniles were less vulnerable to predation. In addition, juvenile blue crab growth rates are higher in upriver unstructured SME due to high availability of preferred prey items (Seitz et al., 2003; Seitz, Lipcius and Seebo, 2005; Lipcius et al., 2005). The shift in utilization from seagrass to SME likely reflects changes in predation risk-growth rate tradeoffs and the changing resource requirements between small and medium-sized juveniles, such that juvenile crabs may derive a growth advantage by dispersing from seagrass beds to SME as they continue to grow (Blackmon and Eggleston, 2001; Reyns and Eggleston, 2004; Reyns, Eggleston and Luettich, 2006; Lipcius et al., 2005; Ralph, 2014; Hyman et al., 2023).

#### 4.1.3. Large juveniles

Large (31–60 mm) juveniles were most abundant in SME. In contrast, large juvenile abundances in seagrass and sand were relatively low and equivalent. Densities in SME were six times as high as those in seagrass and sand. The findings that both medium-sized and large juveniles remain in SME is supported by movement patterns across a range of juvenile sizes in salt marsh habitat, which indicates immature blue crabs between 20–60 mm exhibit high site fidelity and low emigration rates within salt marsh tidal creeks (Johnson and Eggleston, 2010). The ontogenetic habitat-shift paradigm (Werner and Gilliam, 1984; Dahlgren and Eggleston, 2000), coupled with literature on blue crab growth and movement within the seascape-nursery concept offer an explanation for these observed patterns. Although the current understanding of ontogenetic blue crab habitat shifts maintains that larger juveniles exploit unstructured bottom habitat for high food availability, unstructured bottom exists both within marsh-fringed tidal creeks and embayments as well as along shorelines devoid of vegetation. Both of these unstructured habitat types harbor high abundances of preferred prey (Seitz et al., 2003; Seitz, Lipcius and Seebo, 2005). Unstructured habitat near salt marshes may be exceptionally productive, harboring higher diversity and abundance of benthic prey relative to comparable unstructured habitat distant from fringing salt marsh vegetation (Seitz et al., 2006). Accordingly, juveniles of many estuarine species commonly utilize multiple unstructured and structured habitat types for foraging and refuge (Nagelkerken et al., 2015). The close proximity between the high-refuge, structurally complex salt marsh vegetation and prey-rich adjacent unstructured muddy bottom confers high survival and growth rates to both medium-sized and large juveniles within salt marshes. Although unstructured bottom distant from marsh habitat likely offers similarly high growth rates, the additional refuge afforded by salt marshes shifts predation risk-growth rate ratios in favor of salt marsh habitat utilization.

### 4.2. Turbidity

Abundance and survival of small and medium-sized juveniles were positively associated with turbidity. This finding agrees with recent evidence detailing positive effects of turbidity over both large (Hyman et al., 2022) and small (Hyman et al., 2023) spatial scales. In addition, the relationship between turbidity and juvenile blue crab abundance was stronger in medium-sized than in small juveniles. The positive association between juvenile blue crab abundance and turbidity may reflect both top-down and bottom-up effects, albeit to varying degrees. First, juvenile blue crabs are positively associated with abundances of preferred prey such as the thin-shelled bivalve *Macoma balthica*, which aggregate in unstructured habitats near estuarine turbidity maxima (Seitz et al., 2003; Seitz, Lipcius and Seebo, 2005). Hence, positive associations between juvenile blue crab abundance and turbidity may be a proxy for the bottom-up effect of benthic food availability. Second, the positive association between turbidity and survival implies that turbidity inhibits foraging efficiency among visually-oriented predators (O’Brien, Slade and Vinyard, 1976; Howson, 2000; Horodysky et al., 2010) which may partially ameliorate predation pressure for juveniles. Although the effect of turbidity on survival was positive, the posterior contained 0 within 80% credible intervals, indicating a relatively high degree of uncertainty. This may be explained by adaptations of estuarine-dependent predators. Although turbidity reduces foraging efficiency of visually-oriented predators, many estuarine-dependent predators rely on chemosensory abilities to forage in low visibility waters (Howson et al., 2022) and are unlikely to experience major impediments to foraging in highly turbid water. Our results indicate that turbidity is most likely a proxy for food availability (i.e. a bottom-up control; Seitz et al., 2003; Lipcius et al., 2005; Seitz, Lipcius and Seebo, 2005), though turbidity may also provide a partial refuge from predation. Moreover, in concert with ontogenetic preferences of larger juveniles for salt marsh habitat, effects of turbidity provide a mechanism for high abundances of 20–40 mm CW juveniles observed in highly turbid salt marsh habitat (Hyman et al., 2022), as this combination of habitat and environmental characteristics would be ideal for minimizing predation risk-growth rate ratios.

### 4.3. Spatiotemporal variation in habitat-specific survival

Within a habitat, juvenile blue crab survival changed across space and time. Seasonally, survival was highest in spring and late fall, and lowest in summer. This curvilinear pattern in survival is consistent with literature on seasonal juvenile blue crab survival rates and corresponds well with patterns in seasonal predation pressure by cannibalistic conspecifics and piscine blue crab predators (Hines and Ruiz, 1995; Rulifson and Dadswell, 1995; Moody, 1994; Hovel and Lipcius, 2001; Facendola, 2010). In salt marsh habitat, both structured and unstructured, survival increased from downriver to midriver and upriver habitats. Spatial differences in survival in juvenile blue crabs have been linked to variation in predator communities along estuarine salinity gradients (Posey et al., 2005), as well as increased alternative prey availability (Seitz, Lipcius and Seebo, 2005; Lipcius et al., 2005). As our results are consistent under either proposed mechanism, additional work is required to determine the extent to which mechanism is driving survival patterns.

In contrast to survival, abundance of juvenile blue crabs did not vary spatially once we accounted for co-varying environmental variables megalopal abundance and turbidity. Both variables exhibited strong spatial gradients across the tributary axis (i.e. downriver to upriver). Hence, spatial variation in juvenile blue crab abundance is likely predominantly controlled by these environmental variables. Although megalopal supply declined precipitously with distance upriver, juvenile blue crab abundances remained high even in the upriver stratum. This highlights the role of density-dependent secondary dispersal (Etherington and Eggleston, 2000; Reyns and Eggleston, 2004; Blackmon and Eggleston, 2001). High initial settlement of juveniles in downriver structured habitats may exceed habitat carrying capacities, such that juveniles emigrate to upriver habitats to avoid adverse density-dependent effects.

### 4.4. Within-habitat variation in survival

Within seagrass and SME, survival varied markedly among structured and unstructured treatments. In all cases, survival was higher in structurally complex vegetation within seagrass and SME. This finding was particularly notable in seagrass, which conferred the highest survival among all habitats considered in its structured treatments and the lowest survival within its unstructured treatment. These patterns are consistent with the general predation-refuge paradigm associated with structurally complex habitats and edge effects (e.g. Peterson et al., 2001; Smith et al., 2011) and highlights a need for researchers to carefully define the degree of spatial separation between vegetated treatments and unstructured controls in survival studies (Lipcius et al., 2007). The higher refuge value of structurally complex habitats makes them attractive for vulnerable prey. Although the structural complexity of seagrass and salt marsh shoots provides high refuge value, it is also attractive to predators due to the higher availability of food resources (Smith et al., 2011). Within structured portions of these habitats, higher predation pressure is offset by superior refugia. However, interstitial patches devoid of vegetation are characterized by intense predation pressure as in adjacent structured patches but without the associated refuge, leading to disproportionately higher mortality. These effects underscore the need to separate unstructured controls and vegetated treatments by considerable distances to avoid confounding predation pressure and refuge capacity (see details in Lipcius et al., 2007).

These findings have additional implications for intertidal habitats such as salt marshes. As SME remains inundated for only a portion of the full tidal cycle, a high proportion of small juveniles must leave the vegetated marsh surface to adjacent unstructured subtidal habitat at low tide (de la Barra et al., 2022), though some fraction of juveniles may bury in the exposed marsh surface (Johnson and Eggleston, 2010) and adults may remain in tidal pools to opportunistically forage (Johnson, 2022). Although survival in adjacent unstructured habitat in these locations maybe be higher than in similar deeper habitat (Ruiz, Hines and Posey, 1993), these areas nevertheless provide lower refuge quality relative to structured habitat (Ryer, van Montfrans and Moody, 1997). Aggregate survival in SME may therefore be an average in survival across structured and unstructured patches of a nursery habitat, based on accessibility dictated by hydrology and tidal regimes.

### 4.5. Ontogenetic habitat shifts for the juvenile blue crab: a revised paradigm

Placed within the context of previous work, our results indicate that the current paradigm of juvenile blue crab ontogenetic habitat utilization requires revision. Here, we propose a revised conceptual model describing differences in juvenile blue crab habitat use at different size classes (Fig. 8). Initially, ingressing megalopae predominantly settle into seagrass beds or other SAV until *∼*15 mm CW (Orth and van Montfrans, 1987; Hyman et al., 2023; Johnston and Lipcius, 2012), although small juveniles may utilize salt marsh vegetation or algal patches as an alternative nursery if they do not encounter downriver SAV as megalopae (Stockhausen and Lipcius, 2003; Wood and Lipcius, 2022) or after emigrating from seagrass to avoid adverse density-dependent effects (Etherington and Eggleston, 2000; Etherington, Eggleston and Stockhausen, 2003; Reyns and Eggleston, 2004; Blackmon and Eggleston, 2001). Once juveniles reach *∼*15 mm CW, they begin dispersing to SME, algal patches and tidal marsh creek habitats with increasing frequency to exploit the predation refuge and higher food availability (i.e. higher growth rates) associated with these habitats. Juveniles may also exploit refugia associated with salt marsh vegetation, particularly when molting (Ryer, van Montfrans and Moody, 1997; Lipcius et al., 2007; Hines, 2007). The higher food availability in upriver salt marsh habitat near the estuarine turbidity maximum appears particularly valuable for juveniles *≥*20 mm (Hyman et al., 2022). By 31-60 mm CW, most juveniles reside in salt marsh habitats before maturing.

**Fig 8.**
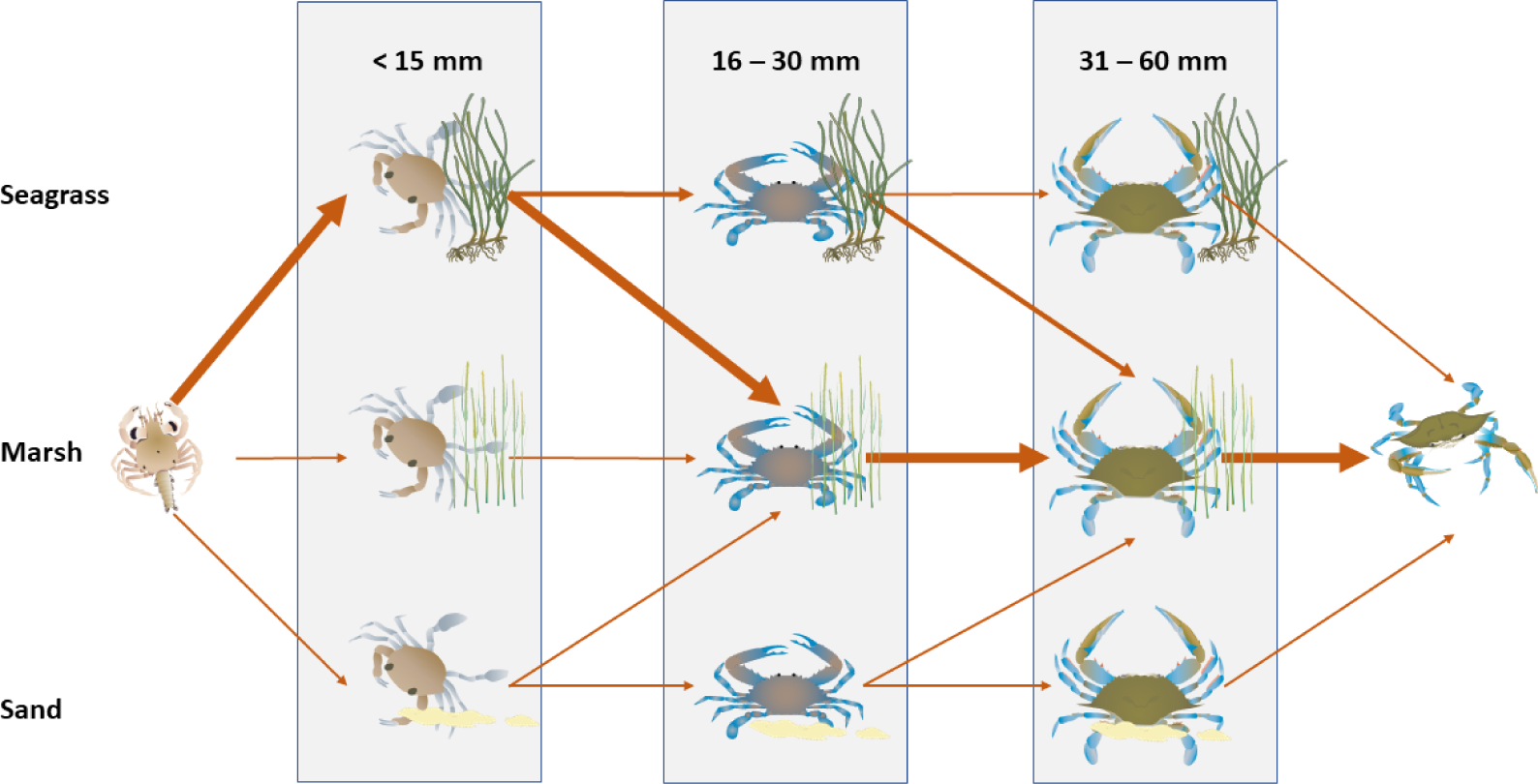
*Conceptual diagram of revised juvenile blue crab ontogenetic habitat shifts. Arrows depict transitions between habitats with increases in size. Arrow widths denote abundance contributions of individuals between habitats*.

Similar ontogenetic habitat shifts were posited by Lipcius et al. (2007), which maintained that the size refuge obtained by juveniles at approximately 30 mm allowed individuals to exploit the high food availability afforded by upriver unstructured habitats. The new conceptual model advanced here illustrates the intermediate refuge value of both unstructured as well as structured salt marsh habitat in conferring high survival and growth rates.

### 4.6. Relevance

Our results expand upon previous work documenting patterns of habitat utilization in juvenile blue crabs to include more size classes and provide a plausible mechanism – trade-offs between growth and survival – underlying ontogenetic habitat shifts. Our findings indicate that shifts in habitat utilization from seagrass beds occur at earlier size intervals than previously thought and emphasize the role of structured salt marsh habitat as a critical nursery for young juveniles.

Our results also add to a growing body of research highlighting a need to protect and restore both seagrass beds and salt marsh habitat to preserve the complete chain of habitats used by juvenile blue crabs before entering adult habitats (e.g. Johnson and Eggleston, 2010; Hyman et al., 2022, 2023). Seagrass and salt marsh declines have received considerable attention in Chesapeake Bay (e.g. Moore, Shields and Parrish, 2014; Schieder, Walters and Kirwan, 2018). Eelgrass (*Zostera marina*) beds are declining due to direct and indirect anthropogenic influences such as land-use change and long-term warming of Chesapeake Bay (Orth et al., 2010; Moore, Shields and Parrish, 2014; Patrick et al., 2018). Similarly, salt marshes have been reduced by coastal development and shoreline hardening (Silliman, Grosholz and Bertness, 2009) as well as sea level rise (Kirwan and Megonigal, 2013; Schieder, Walters and Kirwan, 2018). Although the loss of blue crab secondary production associated with seagrass declines has received considerable attention (e.g. Hovel and Lipcius, 2001, 2002; Mizerek, Regan and Hovel, 2011; Ralph et al., 2013; Johnston and Lipcius, 2012), effects of salt marsh loss on blue crab population dynamics remains a major data gap for Chesapeake Bay and other mid-Atlantic estuaries. Moreover, ratios of marsh-migration to marsh-erosion associated with sea level rise are spatiotemporally variable, and higher rates of salt marsh loss are expected in upriver regions such as within the York River (Schieder, Walters and Kirwan, 2018), which are potentially the most valuable for larger juvenile blue crabs. Additional empirical and mechanistic modeling is required to estimate how shifting seagrass and salt marsh distributions, as well as novel nursery habitats such as algal patches, will affect blue crab population dynamics both within Chesapeake Bay and throughout its range generally.

## APPENDIX A: MORE ON PREDICTOR VARIABLES

### A.1. Predictor justification

#### A.1.1. Habitat

Inference on nursery habitat quality with respect to juvenile blue crab abundance and survival was the over-arching objective of this study. Structurally complex habitats harbor higher densities of juvenile blue crabs relative to unstructured habitats due to higher food availability and superior refuge quality (Orth and van Montfrans, 1987; Lipcius et al., 2005, 2007; Johnston and Lipcius, 2012; Ralph et al., 2013; Hyman et al., 2022, 2023). Hence, relative to sand, we expected higher juvenile blue crab densities in seagrass and SME (Orth and van Montfrans, 1987; Heck Jr, Coen and Morgan, 2001; Johnson and Eggleston, 2010; Etherington and Eggleston, 2000; Hyman et al., 2022, 2023). Moreover structurally complex habitats provide higher refuge quality to small prey through the substantial number of interstitial spaces between biogenic structures such as shoots and rhizomes (Hovel and Lipcius, 2001; Orth and van Montfrans, 2002; Johnston and Lipcius, 2012; Miller et al., 2023). We therefore expected higher survival in structurally complex seagrass and SME relative to sand (Long, Sellers and Hines, 2013; Ajemian, Sohel and Mattila, 2015; Bromilow and Lipcius, 2017). However, not all structure provides equally beneficial refuge. For example, smaller interstitial spaces of seagrass shoots may provide superior refuge quality to juveniles relative to the larger spaces characteristic of salt marsh shoots (Orth and van Montfrans, 2002). In addition, most habitats encompass areas both with and without structural complexity or otherwise spatially vary in the degree of structure they afford (e.g. patchiness; Hovel and Lipcius, 2001, 2002; Lipcius et al., 2005). As a result, in the survival model we considered “structured” and “unstructured” treatments in each habitat (see section 2.3.3 for details). We expected high survival in structured components of seagrass and SME and comparatively low survival in adjacent, unstructured portions.

#### A.1.2. Turbidity

Multiple studies have noted positive correlations between blue crab abundance and turbidity (Hyman et al., 2022, 2023). Two potential mechanisms may engender these observed patterns. First, high turbidity is associated with increased juvenile abundance through both bottom-up controls (Seitz et al., 2003). The thin-shelled Baltic clam *Macoma baltica* is a preferred prey item of juvenile blue crabs (Seitz et al., 2003; Seitz, Lipcius and Seebo, 2005) which aggregates near estuarine turbidity maxima and may attract juveniles (Seitz et al., 2003; Seitz, Lipcius and Seebo, 2005; Lipcius et al., 2005). Second, turbidity may provide juvenile blue crabs with protection from visual predators (Cyrus and Blaber, 1987; Marley et al., 2020) and from predation (O’Brien, Slade and Vinyard, 1976; Howson, 2000; Horodysky et al., 2010) through a reduction in light intensity. Upriver unstructured habitat is turbid, whereas similar habitat downriver has lower turbidity, such that upriver unstructured habitat can also serve as an effective nursery (Lipcius et al., 2005; Seitz, Lipcius and Seebo, 2005). Observed patterns between juvenile blue crab abundance and turbidity at regional scales may be a proxy for patterns between juvenile and potentially top-down (O’Brien, Slade and Vinyard, 1976; Ajemian, Sohel and Mattila, 2015) mechanisms (see methods in Hyman et al., 2022, for more details). Hence, we expected turbidity would be positively associated with juvenile blue crab abundance and survival.

#### A.1.3. Stratum

Aside from turbidity, abundance may be influenced by spatial position through spatially correlated, unobserved variables. Hence, stratum was included in our abundance model to avoid confounding variation as well as to assess the effect of spatial orientation. Habitats in different spatial locations along the river axis may harbor different predator guilds, such that inference on impacts of structure among habitats may be confounded by unmeasured effects of differing predator communities and abundances (i.e. predation pressure; Posey et al., 2005; Lipcius et al., 2005). Thus, we included stratum as a categorical fixed-effect in our abundance model and a habitat-stratum variable (i.e. an interaction) in our survival model.

#### A.1.4. Megalopae

Juvenile abundance is initially dictated by megalopae supply (Epifanio, 2007). Although post-settlement dynamics of early juvenile blue crabs are strongly density-dependent at local scales (Etherington, Eggleston and Stockhausen, 2003; Reyns and Eggleston, 2004), early juvenile abundances may be limited by megalopae supply when juvenile populations are relatively low (Etherington, Eggleston and Stockhausen, 2003; Heck Jr, Coen and Morgan, 2001; Heck Jr and Spitzer, 2001). Megalopae abundance was not expected to affect juvenile blue crab survival, and thus was not included as a predictor in our survival model.

#### A.1.5. Carapace width and size class

Juvenile blue crab habitat utilization changes through ontogeny (Lipcius et al., 2005, 2007; Hyman et al., 2023). Smaller juveniles are vulnerable to a larger suite of predators than larger juveniles which achieve a size refuge from smaller foraging species (Lipcius et al., 2005). We therefore included carapace width as a continuous covariate in our survival model. Carapace width was included as a predictor in our abundance model as three separate size classes as response variables.

#### A.1.6. Month

Juvenile blue crab survival fluctuates seasonally. Juvenile blue crab survival is highest in late spring and late fall months, and lowest in summer (Hines and Ruiz, 1995). Seasonal variation in survival is likely influenced by seasonal predator abundances. For example, red drum *Sciaenops ocellatus*, and striped bass *Morone saxatilis* both consume juvenile blue crabs at high rates (e.g. Mosca III, Rudershausen and Lipcius, 1995; Hines, 2007; Lipcius et al., 2007). Abundances of these species fluctuate seasonally in the North Atlantic estuaries as animals utilize estuaries in during early life stages and spawning in summer and early fall (Facendola, 2010; Rulifson and Dadswell, 1995). Hence, we included month as a categorical variable in the survival model to capture seasonality in the form of a step function. In contrast, juvenile blue crab abundances were sampled only within the fall recruitment period and exploratory data analysis did not indicate substantial fluctuations with month or trip. As a result, month was only included in the survival model.

### A.2. Statistical treatment of continuous variables

Both megalopae abundance and turbidity values were transformed prior to inclusion in abundance models. Megalopae in a given location must enter the river through the mouth, and as a result megalopae abundances generally decline with distance upriver as megalopae encounter suitable habitat and settle out of suspension (Stockhausen and Lipcius, 2003). We expected a negative log-linear relationship between local juvenile blue crab abundance and megalopae abundance (Stockhausen and Lipcius, 2003), where increases in megalopae abundance at low levels have larger positive effects on juvenile blue crab abundance than at higher levels, as post-settlement processes dominate when megalopae supply exceeds the carrying capacity of a system (Heck Jr, Coen and Morgan, 2001; Spitzer, Heck and Valentine, 2003; Lipcius et al., 2007). To avoid infinite values when megalopae were not observed, the natural log of megalopae plus a constant (*M* = ln(Megalopae + 1)) was used in lieu of the raw variable. Moreover, as new recruits require time to grow to a size which could be detected by our gear, we used ln(Megalopae+ 1) abundances lagged by four weeks as a predictor for local juvenile abundance.

Similarly, a natural log transformation was applied to turbidity measurements. Log-turbidity was defined as the natural log transformation of Secchi-disk depth, multiplied by -1 (*T* = *−* ln Secchi). The natural log transformation was applied based on the understanding that a threshold exists in water transparency. Assuming that effects of turbidity on juvenile abundance reflect refuge from visually oriented predators (top-down control), small changes in water transparency when water is relatively clear are not expected to substantially affect juvenile abundance as much as small changes in water transparency when water is cloudy (e.g. predation rates by Summer Flounder on mysid shrimp; Howson, 2000). Alternatively, if associations between juvenile abundance and turbidity are related to elevated food availability near the estuarine turbidity maximum, juveniles would presumably remain more sensitive to fluctuations in turbidity at high values compared to clearer waters. Multiplying the variable by -1 facilitates inference on turbidity, instead of water transparency (inverse).

## APPENDIX B: TETHERING METHODOLOGY DETAILS

Tethering involved attaching 30 cm monofilament fishing line to the crab’s carapace with cyanoacrylate glue. The other end of the line was tied to a stake pushed into the sediment; the stake was tied to a location pole 1 m from the crab to minimize effects of artificial structures that could attract predators to the tethered crab. Tethered crabs were allowed to acclimate in laboratory aquaria for 24 h prior to placement in the field.

Prior to field experiments, pilot experiments were used to determine probability of escape (i.e. crabs un-tethering themselves) and potential changes in behavior. In April (i.e. when predation is negligible; e.g. Ruiz, Hines and Posey, 1993; Moody, 1994) five crabs were tethered in two locations (n = 10) and checked daily. In the absence of predation, juvenile crabs remained on tethers for over a week and were able to bury themselves in sediment. We replicated this procedure in lab conditions with 10 additional crabs observed daily for 10 d. Here, only one crab escaped its tether on day eight. Since crabs in our field experiments were deployed for approximately 24 h, we concluded that the number of crabs removed due to escape was negligible.

Predation is evident by a missing crab and either pieces of carapace remaining on the line, chewed pieces of tape and monofilament line, or cut monofilament lines (e.g., Pile et al., 1996; Hovel and Lipcius, 2002; Lipcius et al., 2005). Although uncommon (i.e. *<* 10% of all instances), some tethers were excluded from analysis because the crab molted and only the exoskeleton remained on the tether. However, this was apparent upon retrieval due to the intact molt remaining on the monofilament line. These features were used to distinguish predation from molting in the field experiments. Molts were subsequently recorded and excluded from analysis.

### B.1. Treatment-specific bias

Tethering can introduce treatment-specific bias in survival (Peterson and Black, 1994). For example, tethered crabs may experience lower survival in structurally complex habitats such as seagrass and SME as a result of entanglement, but would not experience the same reduction in survival in sand, such that relative survival rates could not be compared between these habitats. Alternatively, variation in escape behaviors (e.g., crypsis in structurally complex habitat vs. fleeing in sand) may also introduce bias. Extensive work from previous studies examining treatment-specific biases of tethering juvenile crabs in seagrass, SME, and sand have not found interactions between tethering and habitat (Pile et al., 1996; Hovel and Lipcius, 2001; Lipcius et al., 2005; Miller et al., 2023), therefore, we assumed there was no treatment-specific bias in our experiments, which used similar tethering methods and habitats as those in previous studies.

## APPENDIX C: SUPPLEMENTARY TABLES

**Table A1.**
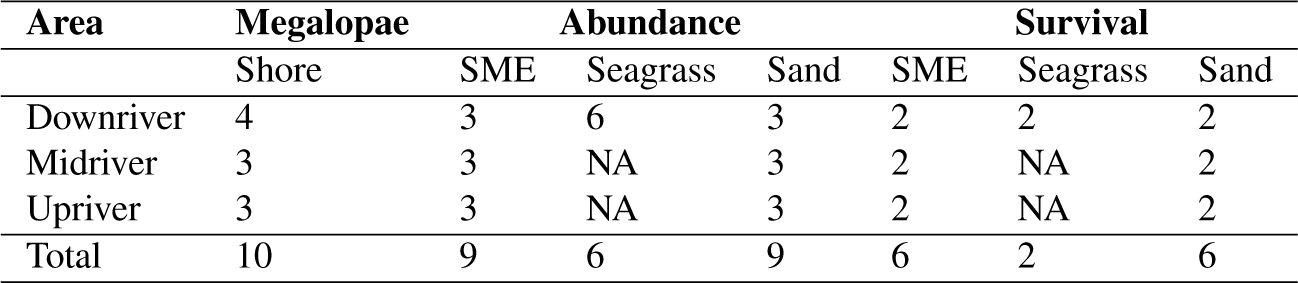
Summary table displaying the number of samples for each habitat by stratum and field study.

**Table A2.** Linear contrast statements depicting differences in juvenile blue crab expected survival (W_j−r_) among habitat-strata combinations. Dots denote posterior median difference in expected values, while thick bars represent 80% Bayesian CIs and thin bars denote 95% Bayesian CIs. The red vertical line denotes 0. Depicted values are on the model (logit) scale. Relevant contrasts are defined as those with their 80% CI excluding 0. Only relevant contrasts are graphed here for brevity.

| Contrast |  | Lower 80% CI | Estimate | Upper 80% CI | Relevant |
| --- | --- | --- | --- | --- | --- |
| Downriver sand | Downriver SME structure | -1.47 | -0.58 | 0.26 |  |
| Downriver sand | Downriver SME unstructured | -0.31 | 0.60 | 1.46 |  |
| Downriver sand | Downriver seagrass structure | -3.07 | -2.19 | -1.32 | * |
| Downriver sand | Downriver seagrass unstructured | 0.40 | 1.37 | 2.27 | * |
| Downriver sand | Midriver sand | -1.27 | -0.32 | 0.57 |  |
| Downriver sand | Midriver SME structure | -2.62 | -1.69 | -0.76 | * |
| Downriver sand | Midriver SME unstructured | -1.27 | -0.35 | 0.57 |  |
| Downriver sand | Upriver sand | -1.24 | -0.24 | 0.68 |  |
| Downriver sand | Upriver SME structure | -3.02 | -2.06 | -1.08 | * |
| Downriver sand | Upriver SME unstructured | -1.33 | -0.38 | 0.56 |  |
| Downriver SME structure | Downriver SME unstructured | 0.70 | 1.19 | 1.65 | * |
| Downriver SME structure | Downriver seagrass structure | -2.41 | -1.61 | -0.80 | * |
| Downriver SME structure | Downriver seagrass unstructured | 1.09 | 1.95 | 2.82 | * |
| Downriver SME structure | Midriver sand | -0.54 | 0.27 | 1.20 |  |
| Downriver SME structure | Midriver SME structure | -1.97 | -1.09 | -0.22 | * |
| Downriver SME structure | Midriver SME unstructured | -0.61 | 0.24 | 1.11 |  |
| Downriver SME structure | Upriver sand | -0.52 | 0.35 | 1.33 |  |
| Downriver SME structure | Upriver SME structure | -2.40 | -1.47 | -0.52 | * |
| Downriver SME structure | Upriver SME unstructured | -0.75 | 0.22 | 1.08 |  |
| Downriver SME unstructured | Downriver seagrass structure | -3.62 | -2.80 | -1.97 | * |
| Downriver SME unstructured | Downriver seagrass unstructured | -0.12 | 0.76 | 1.66 |  |
| Downriver SME unstructured | Midriver sand | -1.81 | -0.92 | -0.02 | * |
| Downriver SME unstructured | Midriver SME structure | -3.16 | -2.29 | -1.37 | * |
| Downriver SME unstructured | Midriver SME unstructured | -1.90 | -0.96 | -0.11 | * |
| Downriver SME unstructured | Upriver sand | -1.83 | -0.85 | 0.09 |  |
| Downriver SME unstructured | Upriver SME structure | -3.65 | -2.66 | -1.71 | * |
| Downriver SME unstructured | Upriver SME unstructured | -1.96 | -0.98 | -0.06 | * |
| Downriver seagrass structure | Downriver seagrass unstructured | 2.97 | 3.56 | 4.09 | * |
| Downriver seagrass structure | Midriver sand | 1.02 | 1.87 | 2.78 | * |
| Downriver seagrass structure | Midriver SME structure | -0.38 | 0.51 | 1.40 |  |
| Downriver seagrass structure | Midriver SME unstructured | 0.95 | 1.84 | 2.73 | * |
| Downriver seagrass structure | Upriver sand | 1.02 | 1.96 | 2.94 | * |
| Downriver seagrass structure | Upriver SME structure | -0.88 | 0.14 | 1.04 |  |
| Downriver seagrass structure | Upriver SME unstructured | 0.85 | 1.82 | 2.72 | * |
| Downriver seagrass unstructured | Midriver sand | -2.63 | -1.68 | -0.76 | * |
| Downriver seagrass unstructured | Midriver SME structure | -4.01 | -3.06 | -2.12 | * |
| Downriver seagrass unstructured | Midriver SME unstructured | -2.62 | -1.72 | -0.74 | * |
| Downriver seagrass unstructured | Upriver sand | -2.64 | -1.61 | -0.66 | * |
| Downriver seagrass unstructured | Upriver SME structure | -4.45 | -3.42 | -2.44 | * |
| Downriver seagrass unstructured | Upriver SME unstructured | -2.78 | -1.74 | -0.81 | * |
| Midriver sand | Midriver SME structure | -2.26 | -1.37 | -0.53 | * |
| Midriver sand | Midriver SME unstructured | -0.89 | -0.04 | 0.83 |  |
| Midriver sand | Upriver sand | -0.80 | 0.08 | 0.98 |  |
| Midriver sand | Upriver SME structure | -2.63 | -1.74 | -0.80 | * |
| Midriver sand | Upriver SME unstructured | -0.96 | -0.06 | 0.81 |  |
| Midriver SME structure | Midriver SME unstructured | 0.83 | 1.33 | 1.82 | * |
| Midriver SME structure | Upriver sand | 0.58 | 1.44 | 2.30 | * |
| Midriver SME structure | Upriver SME structure | -1.24 | -0.37 | 0.53 |  |
| Midriver SME structure | Upriver SME unstructured | 0.48 | 1.31 | 2.18 | * |
| Midriver SME unstructured | Upriver sand | -0.74 | 0.11 | 0.95 |  |
| Midriver SME unstructured | Upriver SME structure | -2.56 | -1.70 | -0.82 | * |
| Midriver SME unstructured | Upriver SME unstructured | -0.86 | -0.02 | 0.82 |  |
| Upriver sand | Upriver SME structure | -2.69 | -1.82 | -0.96 | * |
| Upriver sand | Upriver SME unstructured | -0.97 | -0.13 | 0.69 |  |
| Upriver SME structure | Upriver SME unstructured | 1.16 | 1.68 | 2.17 | * |

## APPENDIX D: SUPPLEMENTARY FIGURES

**Fig A1.**
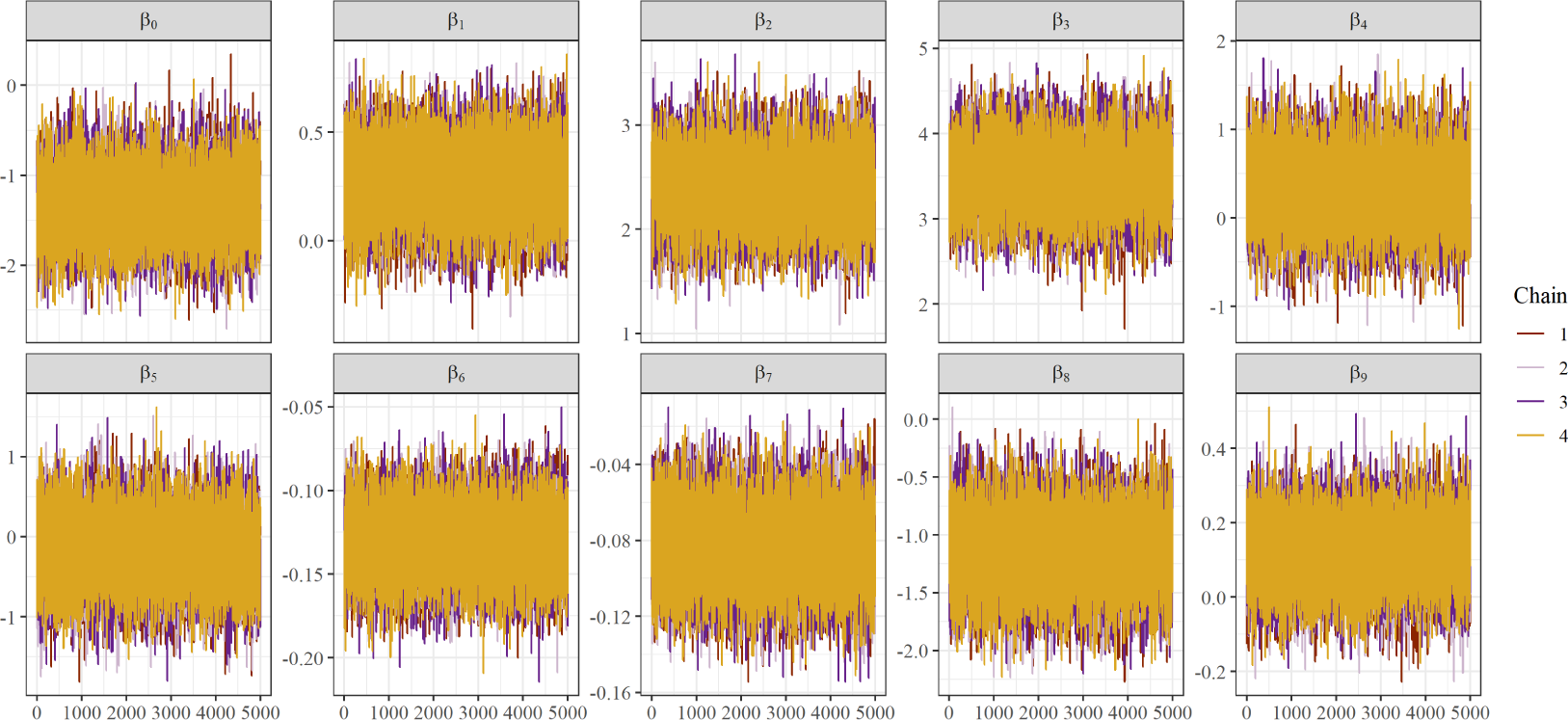
A set of trace plots for abundance model regression parameters for small (≤15 mm) size class illustrating sampled values of regression coefficients per chain throughout the post-warmup/adaptive phase iterations. Visual inspection of trace plots is used to evaluate convergence and mixing of the chains. See *Table 3* for details on abundance model predictor coefficients.

**Fig A2.**
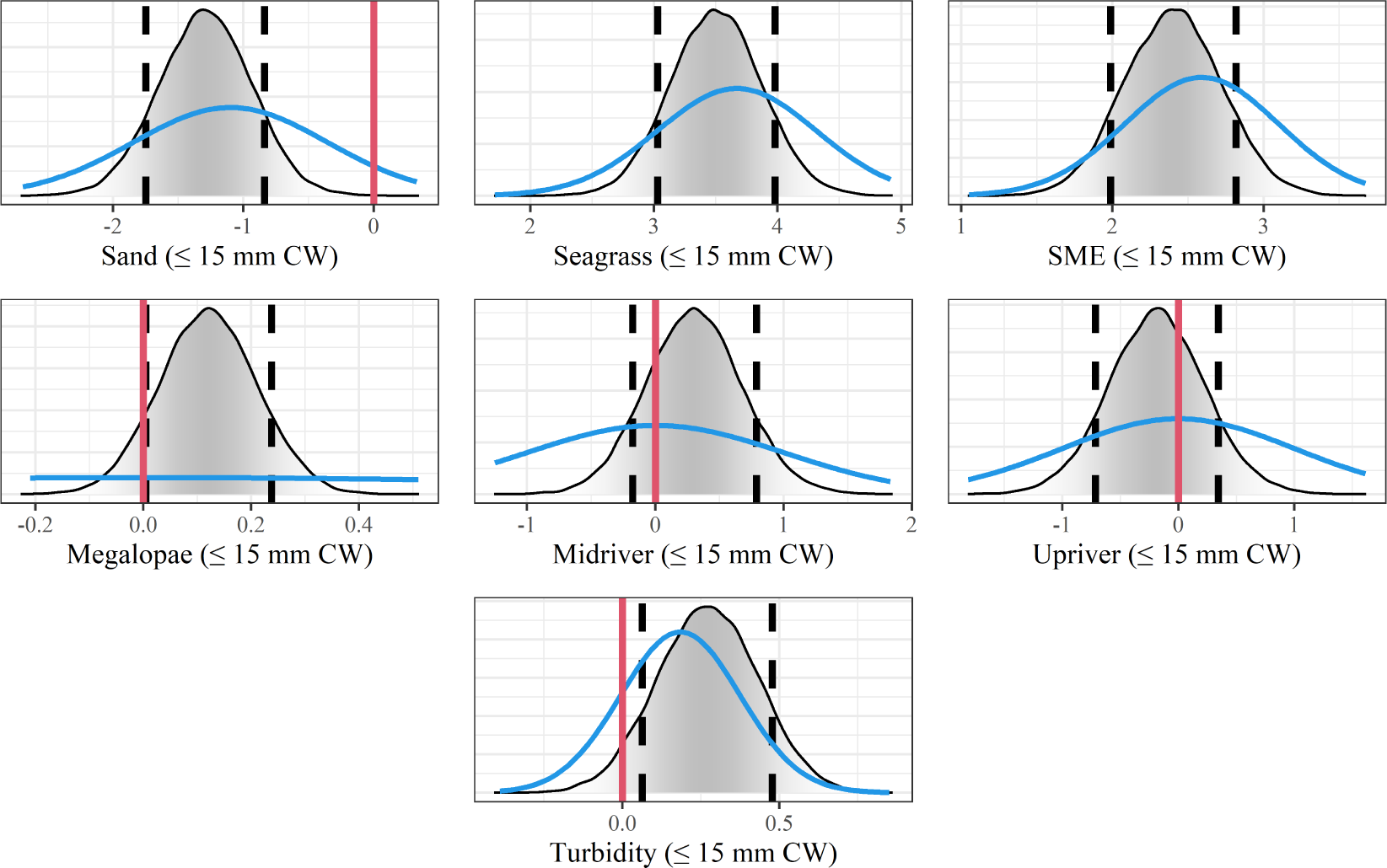
Posterior distributions (grey) and prior distributions (blue) of regression coefficients for small (≤15 mm CW) juvenile blue crabs derived from the abundance model; dashed black lines denote 80% credible intervals, while solid red lines denote 0.

**Fig A3.**
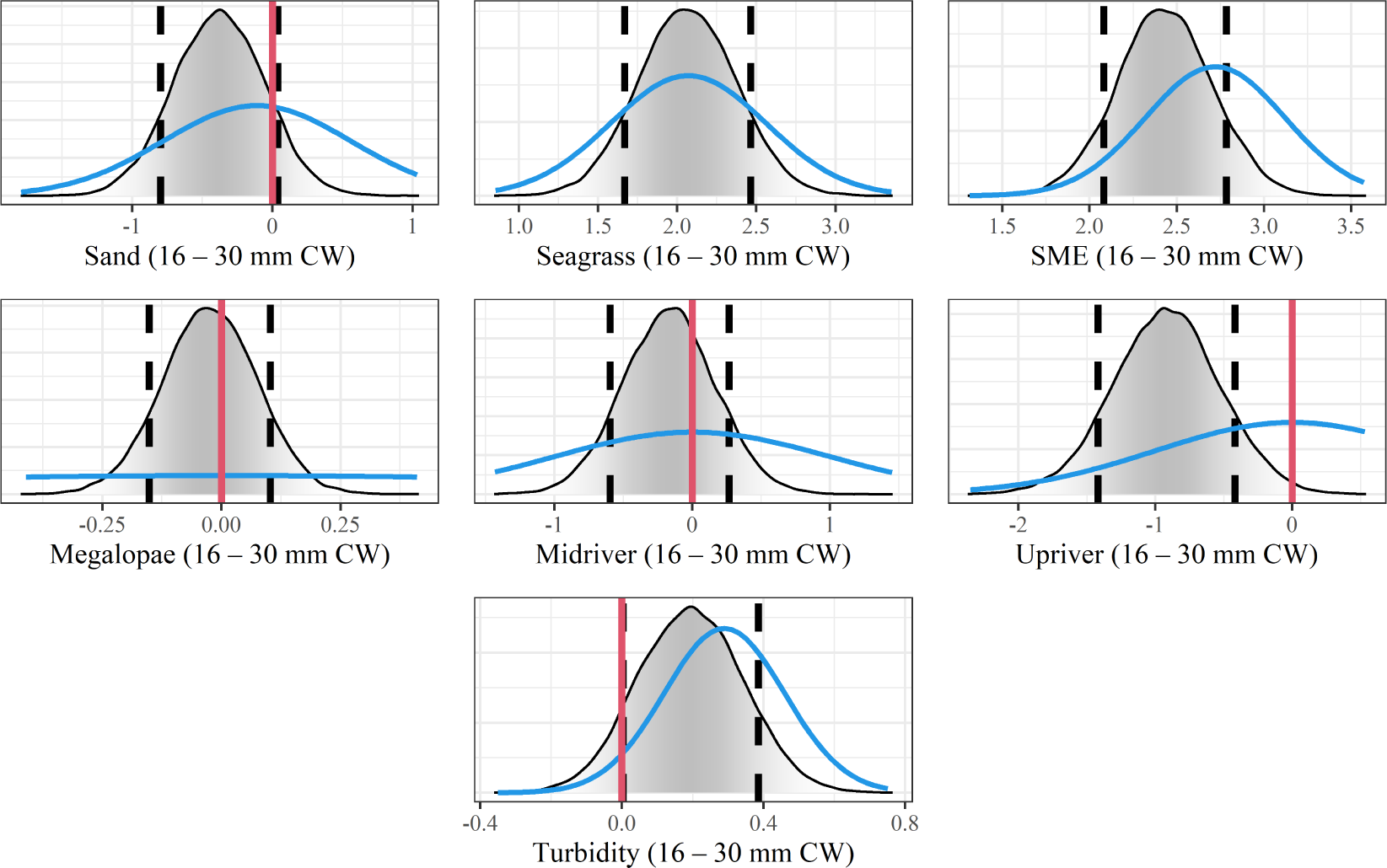
Posterior distributions (grey) and prior distributions (blue) of regression coefficients for medium (16–30 mm CW) juvenile blue crabs in the abundance model; dashed black lines denote 80% credible intervals, while solid red lines denote 0.

**Fig A4.**
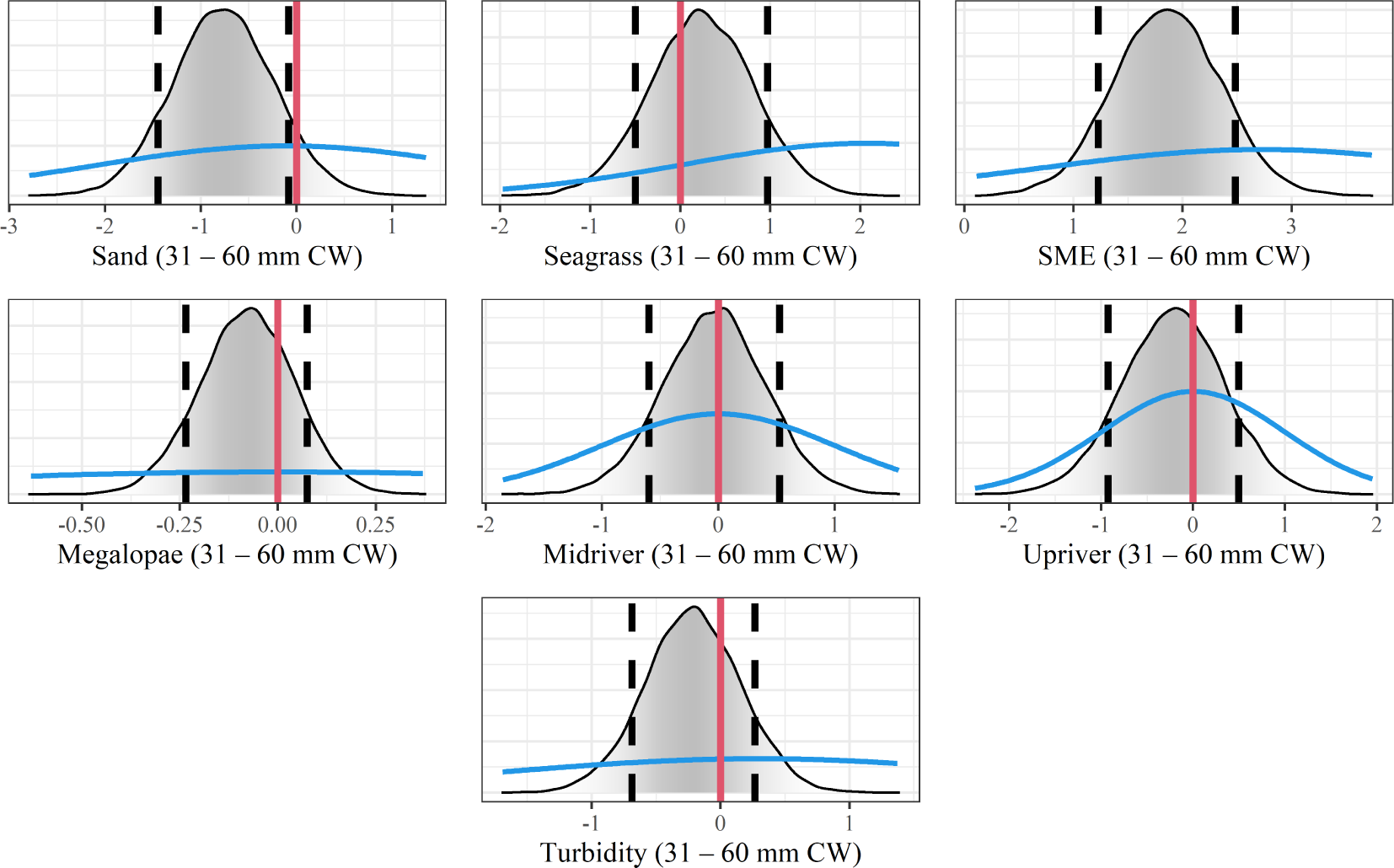
Posterior distributions (grey) and prior distributions (blue) of regression coefficients for large (31–60 mm CW) juvenile blue crabs in the abundance model; dashed black lines denote 80% credible intervals, while solid red lines denote 0.

**Fig A5.**
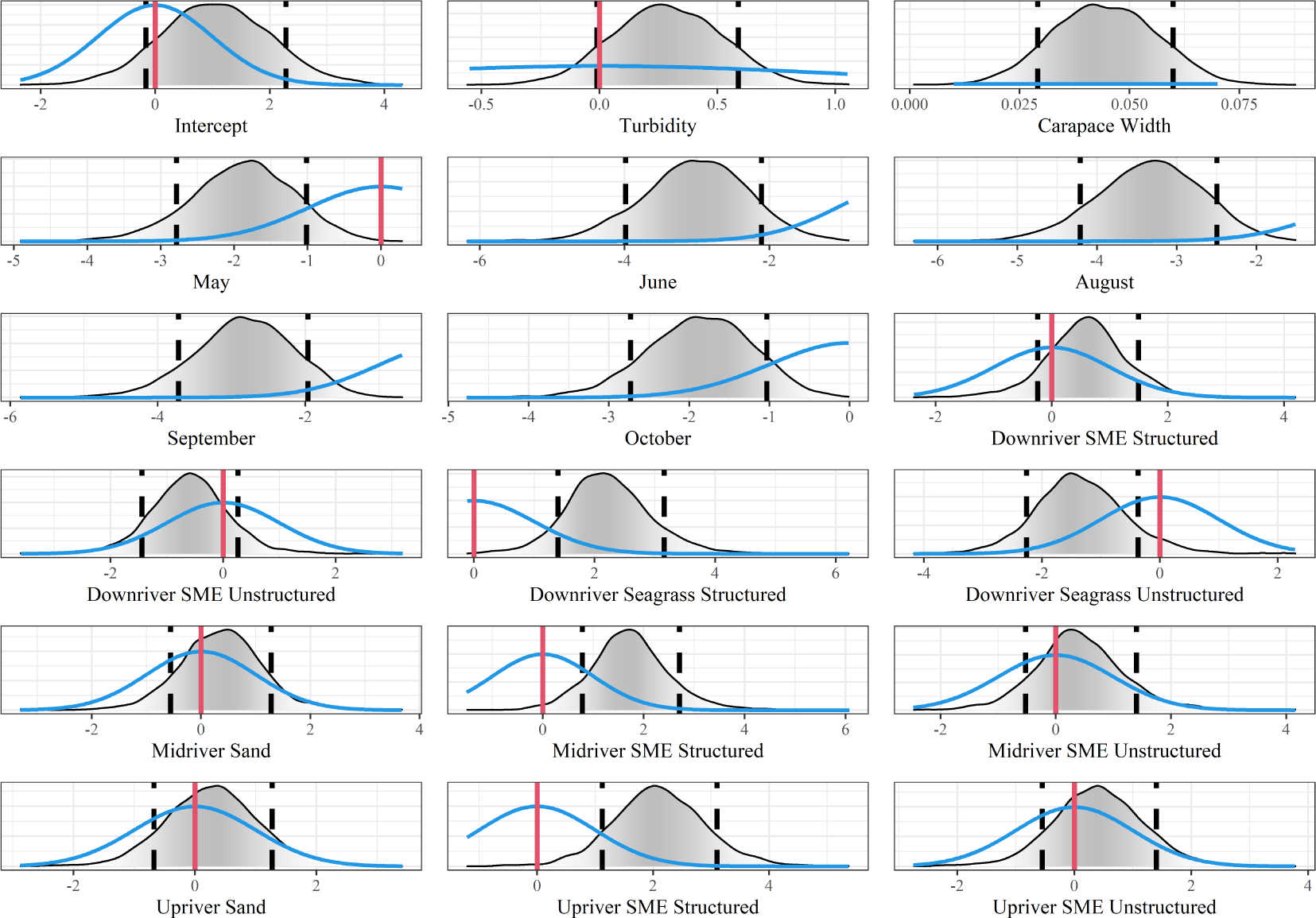
Posterior distributions (grey) and prior distributions (blue) of regression coefficients in the survival model; dashed black lines denote 80% credible intervals, while solid red lines denote 0.

**Fig A6.**
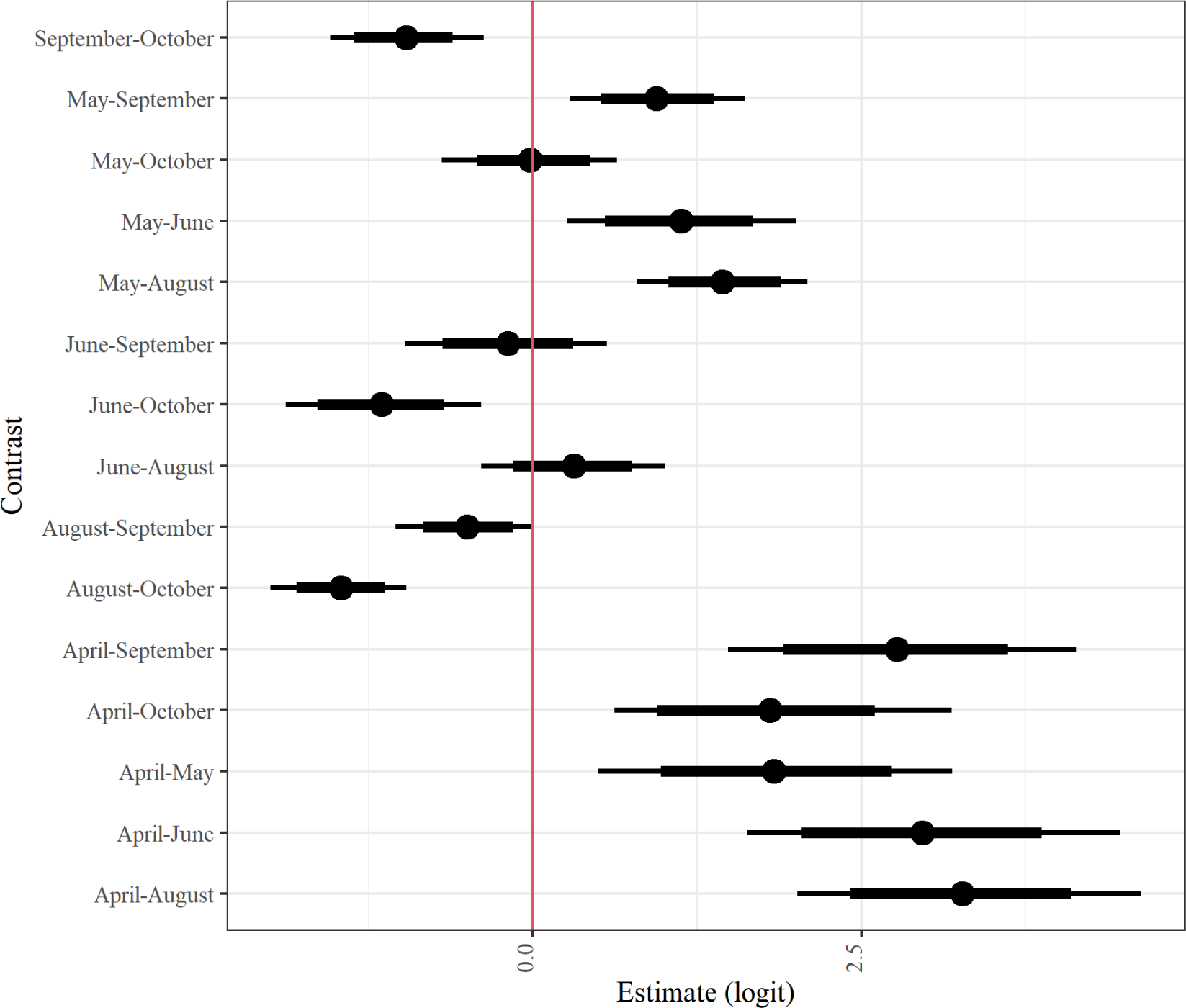
Linear contrast statements depicting differences in juvenile blue crab expected survival among months (W_j−r_). Dots denote posterior median difference in expected values, while thick bars represent 80% Bayesian CIs and thin bars denote 95% Bayesian CIs. The red vertical line denotes 0.

### Acknowledgments

The authors acknowledge William & Mary Research Computing for providing computational resources and technical support that have contributed to the results reported within this paper. URL: https://www.wm.edu/it/rc. ACH also thanks D Eggleston and C Patrick for their ideas as members of ACH’s PhD Committee. Finally, ACH thanks J shields for his guidance and preliminary revisions on this manuscript.

### Funding

Preparation of this manuscript by ACH was funded by a Willard A. Van Engel Fellowship of the Virginia Institute of Marine Science, William & Mary, the NMFS-Sea Grant Joint Fellowship 2021 Program in Population and Ecosystem Dynamics.

